# Structural insights into a conserved mechanism of choline translocation through CHT

**DOI:** 10.1101/2025.09.30.678764

**Authors:** Jesus Vilchez-Garcia, Adrián Martínez-Jiménez, Hanxing Jiang, Borja Ochoa-Lizarralde, Jorge Pedro López-Alonso, Jerónimo Pérez-Lorente, Paola Bartoccioni, Raúl Estévez, Victor Guallar, Ekaitz Errasti-Murugarren, Iban Ubarretxena-Belandia, Igor Tascón

## Abstract

The essential nutrient choline is critical for cellular homeostasis across all domains of life. In humans, choline uptake in cholinergic neurons for its recycling into acetylcholine is mediated by the high-affinity Na⁺-dependent transporter SLC5A7 (also known as CHT1). Prokaryotes also depend on choline as an osmo-protectant and as metabolite, raising the possibility that bacteria also possess choline transporters akin to CHT1. Here, we present a bacterial Na^+^-dependent choline transporter (sfCHT) with high sequence identity to CHT1. sfCHT transport activity can be blocked by the choline transport inhibitor hemicholinium-3. Cryo-EM structures of Na^+^- and choline-bound sfCHT reveal a 14 transmembrane helix topology with a LeuT-fold architecture and Na^+^ coordination geometry similar to CHT1. Captured in an inward-facing conformation, in sfCHT choline is found at a site near the cytoplasmic side. Computational analysis and transport assays reveal local conformational changes along a choline translocation pathway to the cytosolic site. Transport assays with CHT1 variants, carrying substitutions at conserved residues along the proposed translocation pathway in sfCHT, reveal a conserved mechanism of choline transport between the bacterial and human choline transporters.

## Introduction

Choline is a water-soluble quaternary ammonium cation that requires active transport to cross cell membranes and plays important structural and metabolic roles in prokaryotic^1^ and eukaryotic cells^2^. Choline is a building block of phospholipids^3,4^ and also a precursor of the neurotransmitter acetylcholine (ACh). Among the different types of choline transporters in humans, the high-affinity Na^+^-dependent choline transporter SLC5A7 (also known as CHT1) stands out for its role in mediating the uptake of choline, required for its recycling into ACh, at the presynaptic membrane of cholinergic neurons^5^. CHT1 is a Na^+^/choline symporter^6–8^ that exploits the electrochemical gradient of Na^+^ ions to drive the transport of choline. Recent high-resolution structures of human CHT1 in the presence of choline, Na^+^, Cl^-^ and the inhibitors hemicholinium-3 (HC-3) and ML352^9–11^, have revealed the canonical LeuT fold architecture of the transporter, the nature of its ion-binding sites, and key aspects of substrate recognition. However, the mechanism of choline translocation remains unresolved.

In addition of being a structural constituent of some prokaryotic cell membranes^4^, bacteria also rely on choline as a precursor of osmo-protectants such as glycine betaine^12^. In prokaryotes, choline can also serve as a sole source of carbon, nitrogen and metabolic energy^1,13^. Choline and its derivatives can modulate the expression of genes involved in virulence, such as those encoding hemolytic phospholipase C and phosphorylcholine phosphatase^14^. While several bacterial choline transporters have been well studied and even structurally characterized^15–20^, a large fraction remain unstudied. Homologous sequences to the choline transporter CHT1 have been identified across bacterial genomes, with some approaching 40% amino acid sequence identity. However, it remains speculative whether these proteins are capable of choline transport or share any structural and mechanistic determinants with CHT1.

Here, we present a CHT from *Salimicrobium flavidum* (sfCHT) capable of Na^+^-dependent choline transport. Na^+^- and choline-bound cryo-EM structures of sfCHT reveal the same LeuT fold as CHT1, and capture the transporter in an inward-facing conformation with choline bound at a site near the cytoplasmic side. Together with computational analysis and choline uptake assays, we uncover the sequential movements of choline from the substrate-binding site toward the cytosol. Transport assays with CHT1 variants, carrying substitutions at conserved residues along the choline translocation pathway, reveal a conserved mechanism of choline translocation between the bacterial and human choline transporters.

## Results

### A prokaryotic Na^+^-dependent choline transporter inhibited by HC-3

We selected sfCHT from the halophilic bacterium *Salimicrobium flavidum* following an expression and purification screening of prokaryotic homologues displaying a 34-38% amino acid sequence identity and a similar predicted topology with CHT1 (**Supplementary Fig. 1**). sfCHT was expressed in *E. coli* as a C-terminal 3C-His_10_-tag fusion, solubilized in dodecyl-β-D-maltoside (DDM), and purified by a combination of Ni-NTA and size exclusion chromatographies (**Supplementary Fig. 2**). Proteoliposomes (sfCHT-PLs), prepared by reconstitution (1:50 protein to lipid ratio) of purified sfCHT in *E. coli* polar lipid extract bilayers (**Supplementary Fig. 2**), displayed robust sodium-dependent [^3^H]-choline accumulation (linear up to 8 min) in their lumen (**Fig. 1a**). Additionally, the high-affinity CHT1 inhibitor HC-3^6^ abolished the sodium-dependent [^3^H]-choline uptake by sfCHT-PLs (**Fig. 1b**). Altogether these data identified sfCHT as a bacterial Na^+^-dependent choline transporter inhibited by HC-3.

**Figure 1.**
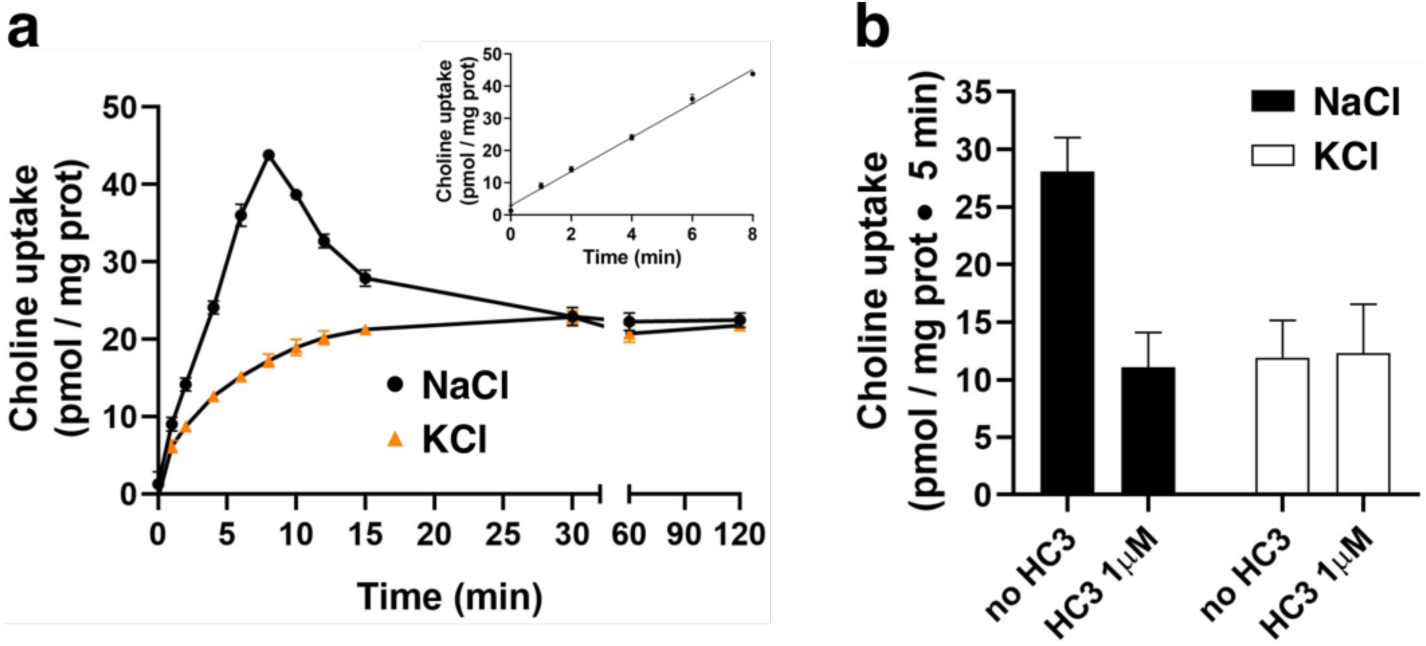
Transport activity of sfCHT reconstituted into *E. coli* polar lipids liposomes. **a)** Time course (0–120 min) of 1 µM [^3^H]-choline (1 µCi/µL) uptake (pmol/mg protein) into sfCHT-PLs in a sodium-containing (137 mM NaCl, black circles) or sodium-free (137 mM KCl, orange triangles) uptake buffer. Inset: Time course (0–8 min) of 1 µM [^3^H]-choline (1 µCi/µL) uptake (pmol/mg protein) into sfCHT-PLs in sodium-containing uptake buffer. Data correspond to a representative experiment, performed using three replicates. A second independent experiment gave similar results. **b)** 1 µM [^3^H]-choline (1 µCi/µL) uptake (pmol/mg protein) into sfCHT-PLs in the presence and absence of 1 μM HC-3 in sodium-containing and sodium-free uptake buffers. Data (mean ± SD) are from three experiments with three replicates per condition.

### LeuT-fold architecture of sfCHT

To elucidate the architecture of sfCHT, we collected cryo-EM images of the transporter in DDM micelles at pH 7.4 in the presence of 100 mM NaCl and in the absence of choline in our in-house 300 kV Krios G4 equipped with a BioContinuum energy filter and a K3 direct electron detector camera. Two-dimensional (2D) classification of the particles extracted from these images yielded a single *ab-initio* class with clear discernible density for transmembrane (TM) helices that remain visible after the detergent micelle region fades out. Three-dimensional (3D) classifications using the good *ab-initio* reconstruction and three decoy models as references followed by non-uniform and local refinements resulted in a 3D-reconstruction at 2.83 Å nominal resolution (**Supplementary Fig. 3, Table 1**).

**Table 1.** Cryo-EM structure determination. Cryo-EM data collection, refinement and validation statistics.

|  | **Na<sup>+</sup>-bound sfCHT**<br>**(EMDB-54347)**<br>**(PDB-9RWT)** | **Choline-bound sfCHT**<br>**(EMDB-54346)**<br>**(PDB-9RWS)** |
| --- | --- | --- |
| **Data collection and processing** |  |  |
| Magnification | 105,000 | 130,000 |
| Voltage (kV) | 300 | 300 |
| Electron exposure (e<sup>-</sup>/Å<sup>2</sup>) | 60.6 | 50 |
| Defocus range (μm) | -1.0 to -1.6 | -1.0 to -1.6 |
| Pixel size (Å/pix) | 0.8238 | 0.6462 |
| Symmetry imposed | C1 | C1 |
| Initial particle images (no.) | 6,671,231 | 5,260,773 |
| Final particle images (no.) | 285,470 | 447,418 |
| Map resolution (Å) | 2.83 | 3.28 |
| FSC threshold | 0.143 | 0.143 |
| Map resolution range (Å) | 2.3 to 4.3 | 2.8 to 4.8 |
| **Refinement** |  |  |
| Initial model used (PDB code) | ModelAngelo model | 9RWT |
| Model resolution (Å) | 2.83 | 3.28 |
| FSC threshold | 0.143 | 0.143 |
| Model resolution range (Å) | 2.3 to 4.3 | 2.8 to 4.8 |
| Map sharpening *B* factor (Å<sup>2</sup>) | -92.6 | -149.7 |
| **Model composition** |  |  |
| Non-hydrogen atoms | 3,895 | 3,903 |
| Protein residues | 512 | 512 |
| Ligands | 1 | 3 |
| ***B* factors (Å<sup>2</sup>)** |  |  |
| Protein | 36.76 | 40.53 |
| Ligand | 24.10 | 62.51 |
| **R.m.s. deviations** |  |  |
| Bond lengths (Å) | 0.004 | 0.002 |
| Bond angles (°) | 0.614 | 0.501 |
| **Validation** |  |  |
| MolProbity score | 1.55 | 1.35 |
| Clashscore | 8.87 | 5.69 |
| Poor rotamers (%) | 0.00 | 0.00 |
| **Ramachandran plot** |  |  |
| Favored (%) | 98.23 | 97.83 |
| Allowed (%) | 1.77 | 2.17 |
| Disallowed (%) | 0.00 | 0.00 |

The cryo-EM map of sfCHT captures the monomeric Na^+^-dependent choline transporter embedded in a DDM micelle (**Fig. 2a**, **Supplementary Fig. 3, Table 1**). The cryo-EM map enabled near-complete *de novo* building of the sfCHT atomic structure (**Fig. 2b, Supplementary Fig. 4**), excluding the first 21 N-terminal and the last 7 C-terminal residues, and the 10 residues in between (70-80) the cytoplasmic loop linking TM helices 2 and 3. The 14 TM helices, with cytoplasmic N- and C-termini, adopt a canonical LeuT fold architecture. The N- (TM helices 3-7) and C-terminal domains (TM helices 8-12) form two inverted structural repeats related by pseudo-two-fold symmetry relative to the membrane plane. These two domains constitute the core of the structure, and are flanked by N- and C-terminal hairpins of two TM helices each oriented toward the cytoplasm (**Fig. 2c**). Overall, the sfCHT structure in the absence of choline resembles the recently determined apo CHT1 structures with a rmsd of 1.057 Å for the 350 matched residues.

**Figure 2.**
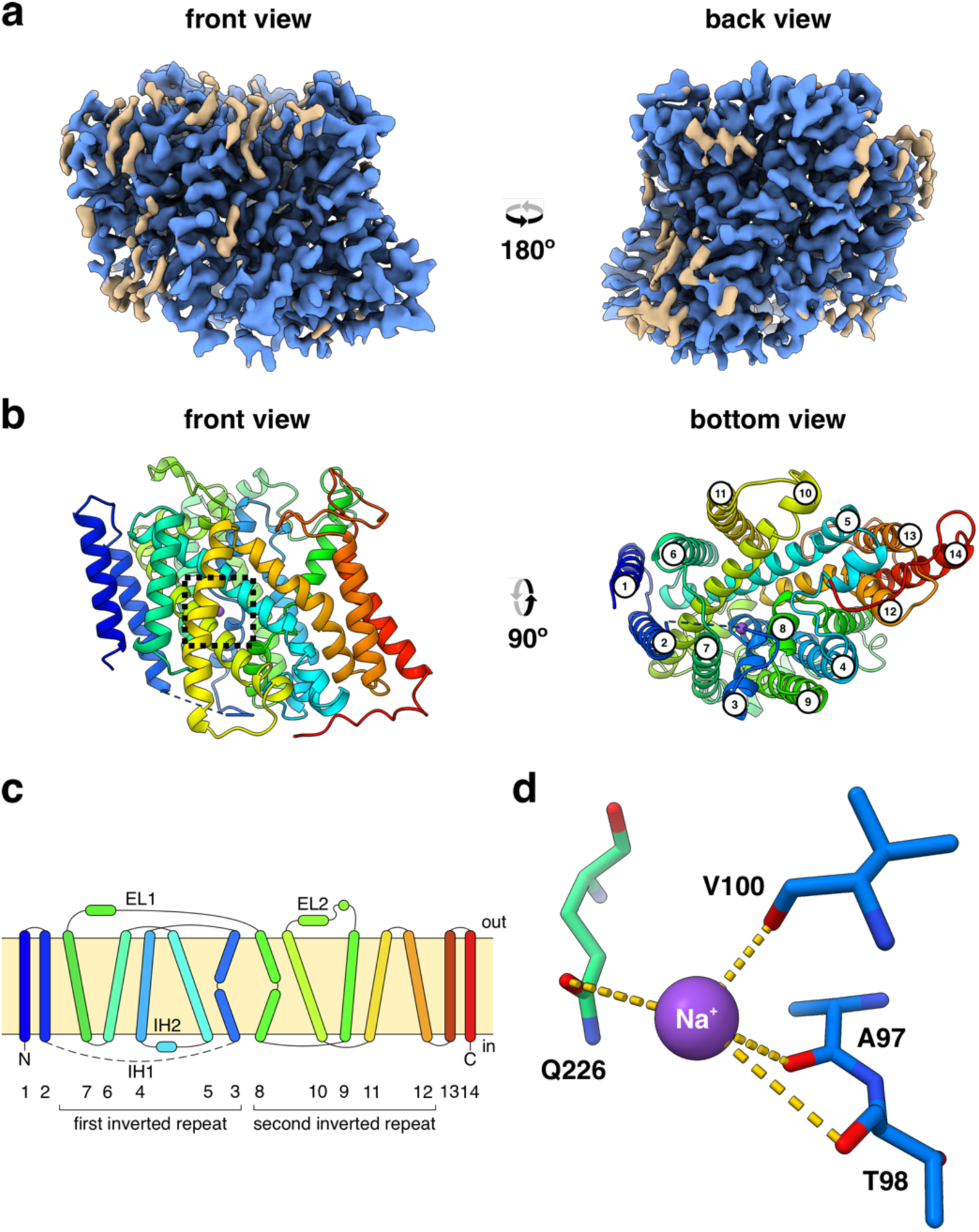
Overview of the structure of Na^+^-bound sfCHT. **a)** Cryo-EM map of sfCHT at 2.83 Å nominal resolution. The front view has been arbitrarily set. The map regions colored in blue correspond to protein and ion-assigned densities, whereas non-protein unassigned densities are brown colored. **b)** Ribbon representation of the atomic model of sfCHT, colored in a rainbow gradient from blue (N-terminus) to red (C-terminus). The missing loop between TM helices 2 and 3 is depicted as a dashed line. Na^+^ cation is shown as a purple sphere. The rectangles in dashed lines highlight the location in the structure of the enlarged view of Na^+^ (d). The numbering of TM helices is shown in the bottom view (right). **c)** Schematic representation of the topology of sfCHT, composed of 14 TM helices, organized in a 5 + 5 inverted structural repeats with two additional TM helices at the N-terminus and two additional TMs at the C-terminus. N, C, IH1, IH2, EL1, EL2 denote the N-terminus, C-terminus, intracellular helix 1, intracellular helix 2, extracellular loop 1, and extracellular loop 2, respectively. The numbering of the TM helices is displayed below, with brackets indicating each of the inverted repeats. **d)** Close-up view of the Na^+^ cation showing the coordination (dotted lines) by interacting residues.

LeuT-fold transporters typically harbor their substrate-binding sites within a central cavity built in between unwound segments of two pseudo-symmetrically related TM helices^21^, which in sfCHT correspond to TM helices 3 and 8. TM helix 3 deviates from canonical α-helical geometry in a region that coincides with the hydrophobic core of the lipid bilayer at residues T98, W99, V100, G101 and G102, dividing the helix in two equally long TM3a and TM3b segments. Similarly, TM helix 8 unwinds at residues G294, G295, I296, P297, W298 and Q299, dividing the helix into a larger TM8a and a shorter TM8b segment (**Supplementary Fig. 5**). The resulting cavity in sfCHT harbors the putative choline-binding site lined by conserved aromatic and polar residues (W99, Y126, W176, W298, W445), which display a similar configuration of that observed in apo CHT1^10^ (**Supplementary 10**). The cavity appears constricted and lacks sufficient space to accommodate a choline molecule. This suggests that conformational rearrangements are required to enable choline binding at the substrate-binding site.

Near the central cavity, a Na^+^ ion could be identified coordinated by residues A97, T98 and V100 in the unwound region of TM helix 3, and by Q226 in TM helix 7 (**Fig. 2d, Supplementary Fig. 4b**). The location of this Na^+^ ion agrees well with the sodium-binding sites observed in other Na^+^-coupled secondary active transporters displaying a LeuT fold, such as the Na^+^/galactose transporter vSGLT (**Supplementary Fig. 7**). Although CHT1 structures in the absence of choline do not display any bound Na^+^ ion, the location matches the observed Na^+^ position in choline-bound CHT1 structures^9–11^.

The structure captures sfCHT in an inward-facing conformation, in which the extracellular side is tightly sealed by packing of TM helices 3b, 4, 5 and 12, while the intracellular side is open to the cytoplasm via an intracellular tunnel (**Supplementary Fig. 8, left**). In contrast, the sfCHT AlphaFold model (AF-A0A1N7IZC0-F1) is closed to both sides of the membrane. We note that in this model the 10-residue cytoplasmic linker connecting TM helices 2 and 3a folds into a short intracellular helix (IH1) that blocks access to the intracellular tunnel from the cytoplasm (**Supplementary Fig. 8, right**). Consistent with the cryo-EM structure representing an inward-facing conformation, in the cryo-EM map of sfCHT this cytoplasmic linker is disordered, which may facilitate the opening of the intracellular tunnel. In the inward-facing structures of CHT1^9–11^ the cytoplasmic loop is also unstructured, suggesting a conserved inward-facing opening mechanism.

### Inward-facing conformation of sfCHT with choline bound at a cytoplasmic site

In an effort to capture the binding mode of choline, we determined the cryo-EM structure of sfCHT at 3.28 Å nominal resolution in the presence of sodium and 1mM choline (**Supplementary Figs. 9 and 10a, Table 1**). The cryo-EM map allowed us to unambiguously build the atomic model for residues 22-69 and 81-543, as in the Na^+^-bound structure (**Supplementary** Fig. 11). Cryo-EM captures sfCHT in the same inward-facing conformation as the Na^+^-bound structure (rmsd of 0.423 Å for the 512 built residues), with a Na^+^ ion found at the same conserved position at the intracellular tunnel near the central cavity comprising the substrate-binding site within TM helices 3 and 8 (**Supplementary Figs. 10c and 11b**). Nevertheless, this site presents a local tightening that allows us to build a water molecule, which was not observed in the Na^+^-bound structure.

We could not identify any density for choline within this cavity, which as in the Na^+^-bound structure was hindered by the side chains of the aromatic and polar residues (W99, Y126, W176, W298, W445). Instead, we found non-protein density consistent with choline close to the cytoplasmic exit of the intracellular tunnel (**Fig. 3a and Supplementary Fig. 11c**). In this site, the trimethylammonium headgroup of choline interacts with the side chain of residues R304, Q303 and S307 from TM helix 8a, while its hydroxyl moiety establishes a hydrogen bond with the backbone amide of residue A220 in TM helix 7 (**Fig. 3a**). The proximity of choline to the cytoplasmic surface of the membrane suggests we may have identified a cytoplasmic site. Consistent with this hypothesis, Q303A, R304A and S307A sfCHT variants displayed a marked reduction in choline uptake activity compared to wild-type (WT) sfCHT in liposome-based transport assays, without affecting either protein expression nor liposome reconstitution efficiency (**Fig. 3c, Supplementary Fig. 12a**). The cytoplasmic site observed in sfCHT has not been reported for CHT1^9–11^.

**Figure 3.**
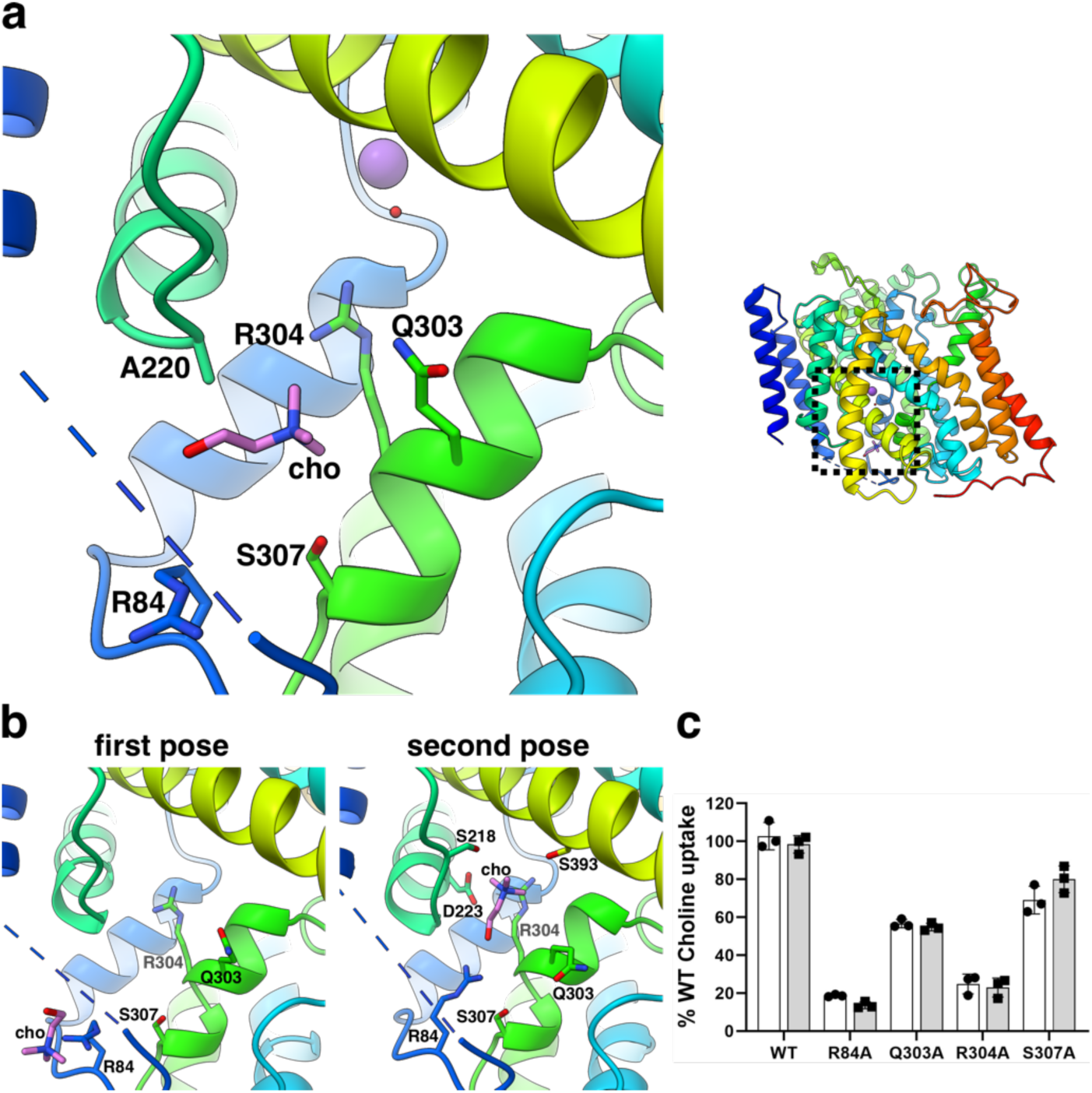
Interaction of choline and sfCHT. **a)** Close-up view of the highlighted region in the experimentally determined choline-bound sfCHT structure, illustrating choline positioned at the cytoplasmic site. **b)** Close-up view of two consecutive choline poses in PELE-predicted models. Residues located within interaction distance of choline and gating residues lining the intracellular tunnel are displayed in sticks. **c)** Transport activity of R84A, Q303A, R304A and S307A sfCHT variants reconstituted in liposomes. 1 µM (white bars) and 50 µM (grey bars) [^3^H]-choline (1 µCi/µL) 5 min uptake into proteoliposomes carrying WT and variant sfCHT is shown. Data are normalized to WT sfCHT choline uptake activity. Data correspond to three experiments with three replicates per condition.

To further assess the relevance of the cytoplasmic site in the mechanism of choline translocation we performed PELE (Protein Energy Landscape Exploration) analysis. The PELE analysis was carried out using the highest resolution cryo-EM structure of sfCHT (Na^+^-bound structure) without any ion and commenced with an unsupervised docking of choline from outside of the transporter to explore different poses. PELE predicted two minimum-energy poses for choline at the exit of the intracellular tunnel, in an arrangement that closely resembles the choline position observed in the cryo-EM map that represents an intermediate between both predicted poses (**Fig. 3b**). Notably, while R84, located in the structured part of the cytoplasmic linker connecting TM helices 2 and 3a, appears in an extended conformation in the second PELE-predicted pose, in the experimentally determined sfCHT structure and in the first PELE-predicted pose, R84 adopts a bent conformation (**Fig. 3a and 3b**). This bent conformation creates space for choline. The substitution R84A in sfCHT resulted in a significantly impaired choline uptake activity, indicating that this residue may contribute to choline translocation (**Fig. 3c, Supplementary Fig. 12a**). We suggest R84 plays a key role in the final step of substrate release to the cytoplasm. Remarkably, in both the cryo-EM structure and the second minimum-energy pose, choline is positioned immediately after Q303 and R304 (**Fig. 3**), implicating these residues as potential gating elements that prevent backflow of the substrate into the intracellular tunnel.

### Structural basis of choline translocation

Starting from the pose at the exit of the intracellular tunnel towards the central cavity, PELE simulations identified additional minimum-energy poses along the tunnel and within the substrate-binding site that help identify the structural determinants underlying choline transit from the substrate-binding site to the cytoplasmic vestibule where choline was observed in the cryo-EM structure. Two intermediate poses of choline within the intracellular tunnel (**Fig. 4a, poses i and ii**) revealed conformational changes of the side chains of residues Q303 and R304, which would permit the passage of choline, supporting their role as gating residues regulating exit from the intracellular tunnel into the cytoplasmic vestibule. Consistently, *in vitro* functional characterization of Q303A and R304A variants resulted in a complete loss of choline uptake (**Fig. 3c, Supplementary Fig. 12a**).

**Figure 4.**
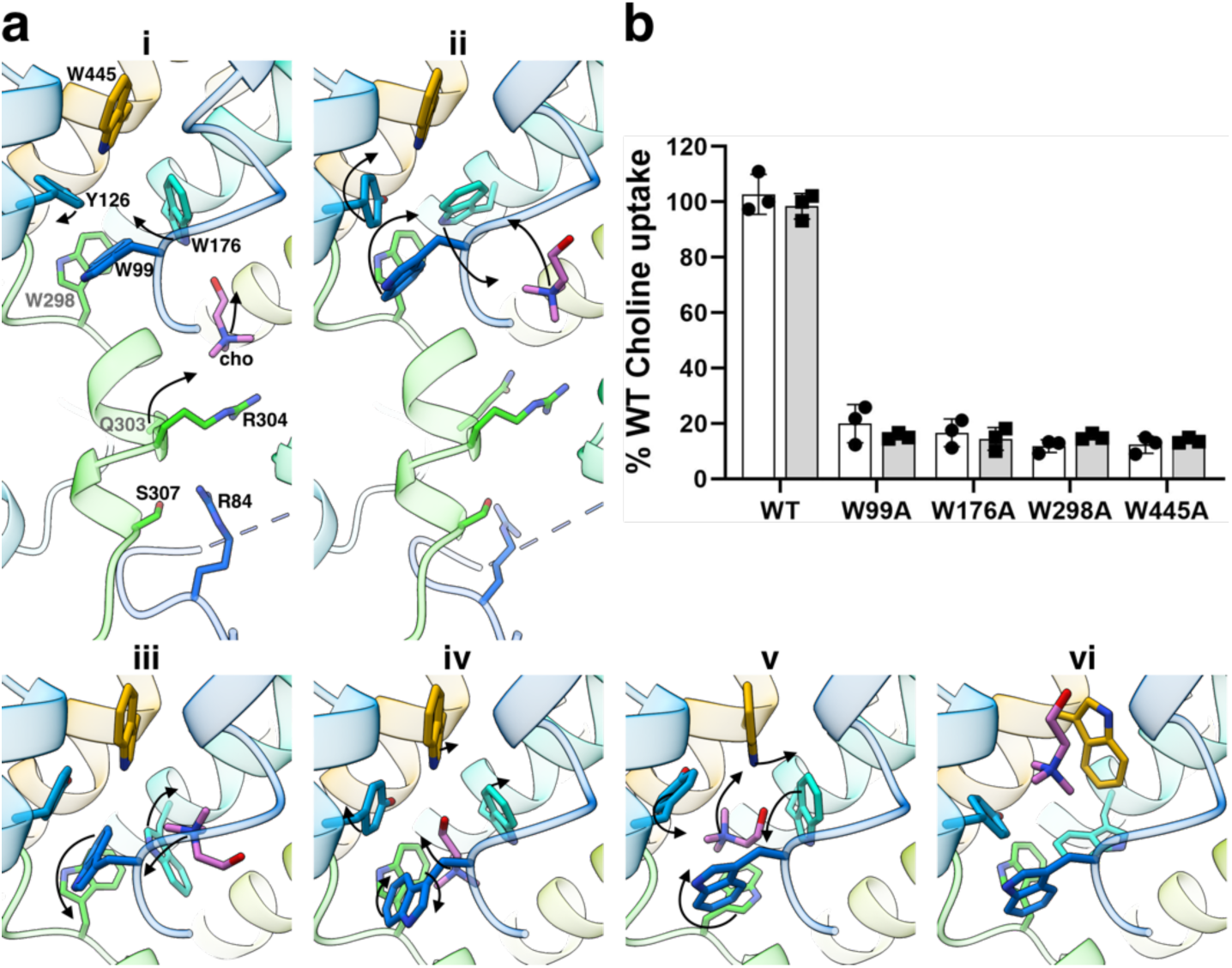
Local conformational changes along the translocation pathway. **a)** Sequential PELE-predicted poses towards the substrate binding site. Panels i and ii provide a view of the translocation pathway, whereas panels iii, iv, v and vi focus on the rearrangements within the substrate-binding site. Side chains displayed as sticks represent PELE-identified key residues in choline-binding and translocation. Arrows indicate the direction of the movement of choline and side chains to the next pose. **b)** Transport activity of W99A, W176A, W298A, W445A sfCHT variants reconstituted in liposomes. 1 µM (white bars) and 50 µM (grey bars) [^3^H]-choline (1 µCi/µL) 5 min uptake into proteoliposomes carrying WT and variant sfCHT is shown. Data are normalized to WT sfCHT choline uptake activity. Data correspond to three experiments with three replicates per condition.

PELE analysis also predicted a minimal-energy pose at the canonical substrate-binding site within the unwound regions of TM helices 3 and 8 (**Fig. 4a, pose v**), as reported for other LeuT-fold transporters, including CHT1^9,10,22–25^. In the predicted substrate-binding site pose, the trimethylammonium headgroup of choline forms ρε-cation interactions with the indole of W176 and W298 in TM helices 5 and 8, respectively, while the substrate hydroxyl group interacts with the imidazole ring of W99 and W445 in TM helices 3 and 12, respectively. Additionally, PELE analysis suggests that conformational flexibility of the side chains of residues W99 and W176 is necessary for choline transition towards the intracellular tunnel (**Fig. 4a, poses iii and iv**), while W445 might also act as an extracellular gate (**Fig. 4a, pose vi**). In support of this, liposome-based transport assays of W99A, W176A, W298A and W445A sfCHT variants showed a total loss of choline uptake activity (**Fig. 4b**). Interestingly, the calculated choline pose observed in our PELE analysis closely resembles that of the substrate-bound conformations observed in CHT1 cryo-EM structures^9–11^, although with significant differences (**Supplementary Fig. 6b**). In CHT1, a Cl^-^ ion is observed within the substrate-binding site where it interacts with cavity-forming residues, accommodating them to optimally coordinate the choline molecule. Whether this Cl^-^ ion may play a role in substrate recognition and/or translocation in CHT transporters needs further investigation. Overall, the coordinated and sequential rearrangement of residues W99, W176, W298, Q303, R304 and R84 appears to create transient pockets that guide choline through the intracellular tunnel (**Supplementary Movie 1**).

### Conserved residues for sfCHT transport activity impact CHT1-mediated choline uptake

To investigate the conservation of choline translocation mechanisms between sfCHT and CHT1, we leveraged the high sequence identity between both transporters (**Fig. 5a, Supplementary Fig. 1**). We carried out a substitution analysis on the conserved residues in CHT1 that show an effect in sfCHT transport activity. Transport assays of CHT1 variants *R47A* (R84)*, Q259A* (Q303)*, R260A* (R304) and *S263A* (S307) (hereby CHT1 numbering italicized and related sfCHT numbering in parentheses) resulted in a nearly identical choline uptake activity profile to that observed for the bacterial counterpart (**Figs. 5b and 4b**), without affecting protein expression (**Supplementary Fig. 12b)**. These findings reinforce a likely conserved key role for the selected residues in the choline translocation mechanism from these two CHT orthologs. In support of this, kinetic characterization of the *Q259A* (Q303) and *S263A* (S307) variants resulted in a reduction of the *V_max_* without affecting choline apparent affinity (**Supplementary Fig. 13, Table 2)**, suggesting a role for these residues in choline translocation rather than recognition.

**Figure 5.**
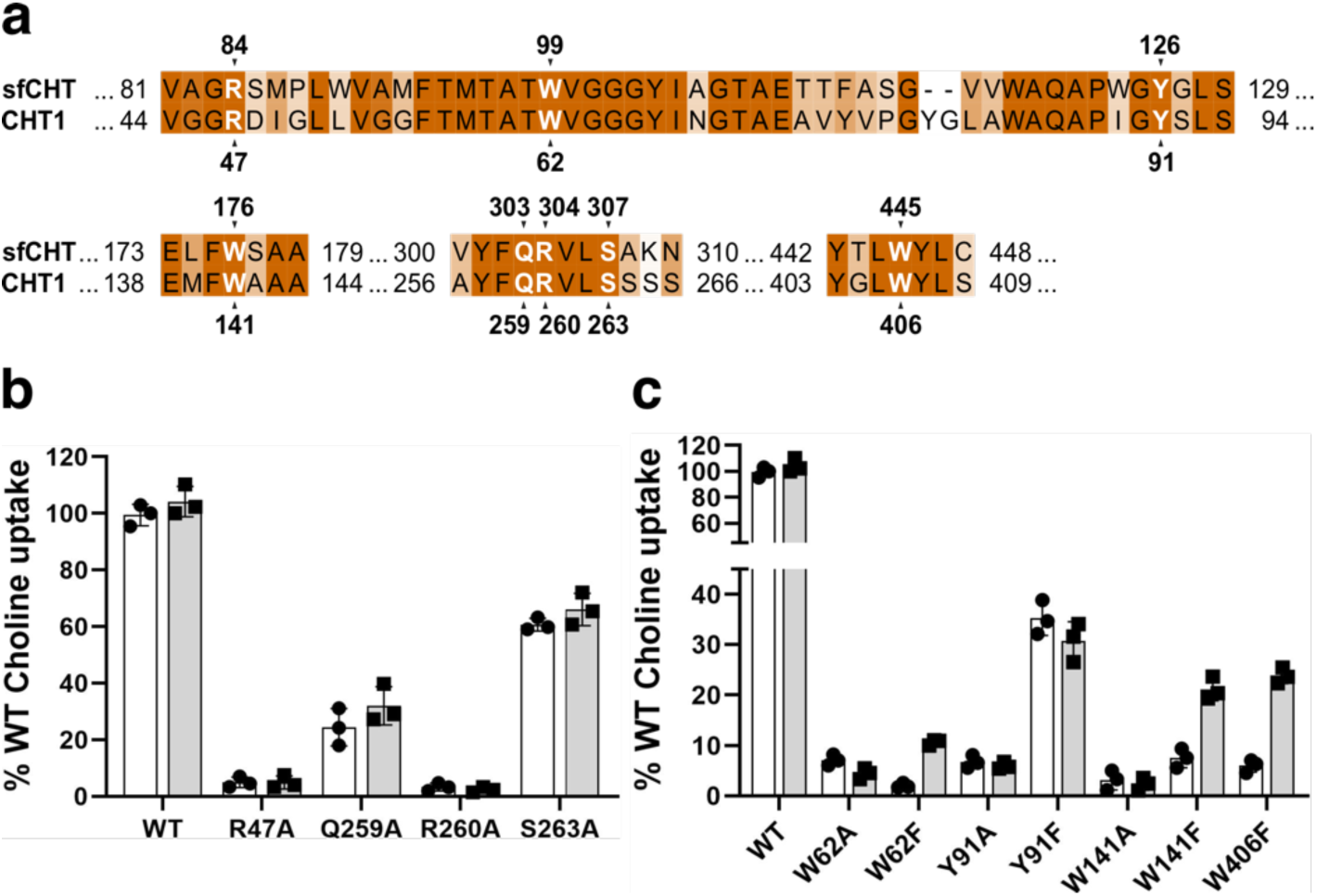
Choline translocation mechanism is conserved in CHT1. **a)** Sequence alignment between sfCHT and CHT1 showing that the key residues in sfCHT, identified by PELE and substitution analysis, are conserved in CHT1. Transport activity of selected CHT1 variants within the translocation pathway **(b)** or located at the substrate-binding site **(c)**. Both panels show 1 µM (white bars) and 50 µM (grey bars) [^3^H]-choline (1 µCi/µL) uptake in HeLa cells by WT and variant CHT1. Uptake activity is normalized for WT CHT1 choline uptake. Data (mean ± SD) corresponds to 3 independent experiments run in quadruplicate. Endocytosis resistant CHT1 LV/AA variant, we refer as WT CHT1 in the context of this study, was used as template for the generation of CHT1 variants.

**Table 2.**
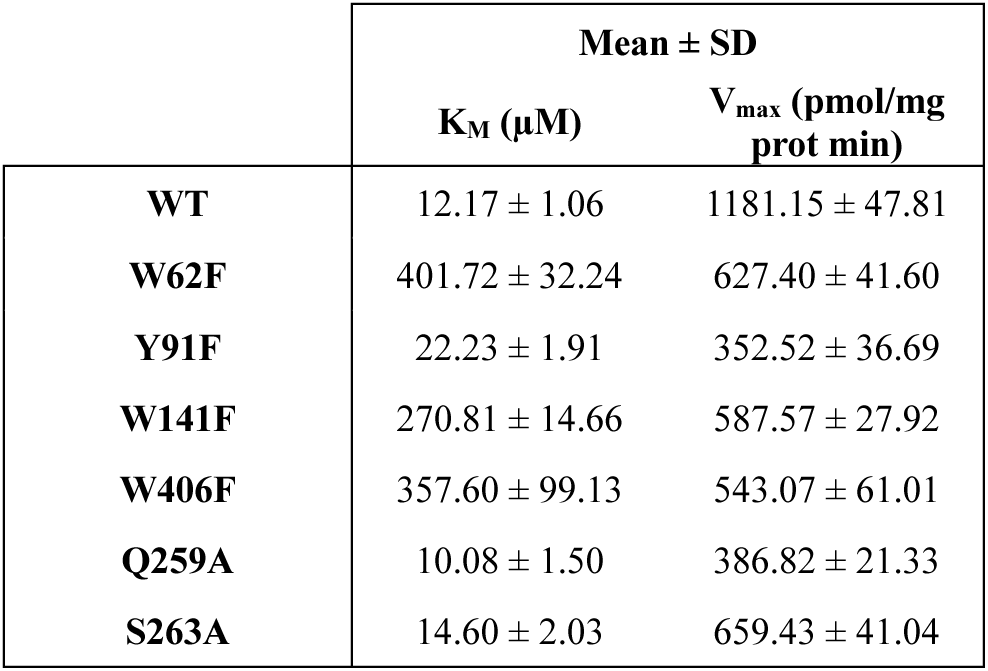
Kinetics parameters K_M_ and V_max_ derived from WT and variant CHT1 uptake of [^3^H]-choline. Data from three representative experiments and mean ± SD are shown.

| | Mean ± SD | |
| --- | --- | --- |
| | $K_M$ ($\mu M$) | $V_{max}$ (pmol/mg prot min) |
| **WT** | 12.17 ± 1.06 | 1181.15 ± 47.81 |
| **W62F** | 401.72 ± 32.24 | 627.40 ± 41.60 |
| **Y91F** | 22.23 ± 1.91 | 352.52 ± 36.69 |
| **W141F** | 270.81 ± 14.66 | 587.57 ± 27.92 |
| **W406F** | 357.60 ± 99.13 | 543.07 ± 61.01 |
| **Q259A** | 10.08 ± 1.50 | 386.82 ± 21.33 |
| **S263A** | 14.60 ± 2.03 | 659.43 ± 41.04 |

Similarly, transport assays of CHT1 variants within the substrate-binding site *W62A* (W99), *Y91A* (Y126) and *W141A* (W176) (**Fig. 5c)**, and *W254A* (W298) and *W406A* (W445)^9,10^, revealed a complete loss of choline uptake activity, with no impact on protein expression (**Supplementary Fig. 12b)**, suggesting a key role for the selected residues in choline recognition and/or translocation mechanisms. Interestingly, substitution of some of the above mentioned residues to phenylalanine resulted in a partial recovery of the choline uptake activity (**Fig. 5c**). Kinetic characterization of *W62F*, *W141F* and *W406F* CHT1 variants resulted in a one order of magnitude reduction in choline apparent affinity, together with a ≈ 50% reduction in *V_max_*. Similarly, *Y91F* CHT1 variant showed a reduced *V_max_* value, although choline apparent affinity was barely affected (**Supplementary Fig. 13, Table 2**). All these observations point towards a key role for the aromatic residues in the choline-binding site, both in substrate recognition and translocation. Indeed, choline-binding site cavity volume increases as a result of tryptophan and tyrosine residues substitution, may result in non-canonical substrate poses, reducing choline apparent affinity, and ultimately, substrate uptake.

Collectively, the results of the residue substitution analysis of CHT1 point towards a mechanism of choline recognition and translocation, from the substrate-binding site to the intracellular space, mirroring that of sfCHT. These findings reveal an evolutionarily conserved choline transport mechanism between sfCHT and CHT1.

## Discussion

Choline is a structural component of prokaryotic cell membranes^4^, and its uptake is a critical adaptive strategy in many bacterial species, enabling survival under osmotic stress^12^, serving as carbon, nitrogen and energy source^1,13^, and acting as a virulence factor^14,19^. In this study, we have identified a prokaryotic Na^+^-dependent choline transporter from *Salimicrobium flavidum* (sfCHT), with high sequence identity to CHT1. Cryo-EM structures of sfCHT in Na^+^- and choline-bound inward-facing conformations reveal a LeuT fold architecture, Na^+^-binding sites similar to LeuT and other sodium-coupled symporters, as reviewed^26^, including CHT1^9–11^, and an unprecedented cytoplasmic choline-binding site. The cryo-EM structures combined with computational and transport assays, provide mechanistic insights into substrate release from the binding site and reveal substrate-induced local conformational changes along the translocation pathway. Comparative substitution analysis between CHT1 and sfCHT highlights the conservation of the choline transport mechanisms and advances our understanding of CHT-mediated choline translocation. While CHT1 plays a well-established role in neuronal choline uptake for acetylcholine biosynthesis, the biological function of its prokaryotic homologs remains largely unexplored. Future investigations will be essential to elucidate these roles, which could reveal novel insights into bacterial biology and potentially identify new targets for antimicrobial strategies.

Secondary active transporters operate via an alternating access mechanism, transitioning between outward- and inward-facing states through intermediate occluded conformations^26,27^. The sfCHT cryo-EM structures capture an inward-facing conformation with a tightly sealed extracellular side and a wide-open intracellular tunnel. In contrast, the AlphaFold model of sfCHT predicts a short helix (IH1) within the linker between TM helices 2 and 3 that blocks cytoplasmic access to the intracellular tunnel, resembling the outward-facing conformations of other SLC5 transporters such as CHT1, SGLT1 and SGLT2^9,10,28^. Notably, IH1 is unstructured in inward-facing CHT1 structures^9–11^, suggesting that during inward transition, the unwinding of IH1 releases the intracellular ends of the TM helices, enabling tunnel formation and substrate passage. These findings support a conserved transition mechanism between outward and inward conformations between sfCHT and CHT1.

The choline-bound sfCHT structure captures a stable intermediate state with choline bound to a substrate-release site (**Fig. 3**), which together with PELE analysis and transport assays of sfCHT key residue variants, provides new clues for understanding the choline translocation mechanism through CHT (**Fig. 6, Supplementary Movie 1**). We posit that in sfCHT the translocation process starts after substrate release from the binding site, facilitated by the rearrangement of the proposed intracellular gate, residues W99 and W176 (**Fig. 4**). Choline is sandwiched between the aromatic residues of the substrate-binding site and Q303 and R304, which functions as an intermediate gate (**Fig. 4**). These long side chain residues might serve a dual role: acting as bulky residues that confine the small choline substrate to prevent premature access to the intracellular vestibule; and as flexible elements which undergo conformational changes to enable choline translocation to proceed. Binding of choline to R84 subsequently induces the closure of the intermediate gate, forming a stable conformational state. Q303 and R304 reconfiguration after substrate release will effectively prevent the substrate back diffusion. Furthermore, the cryo-EM determined intermediate state reveals R84 in a bent conformation, representing a stage preceding choline release from the transporter vestibule into the intracellular space. Additionally, S307 that coordinates choline in the cryo-EM structure, appears to contribute to translocation but it is not essential, consistent with its limited conservation among bacterial homologs. These findings entail a more detailed, stepwise mechanism for choline translocation through sfCHT (**Fig. 6, Supplementary Movie 1**). This dynamic process may ensure the translocation of the small substrate choline from the canonical substrate-binding site to the cytoplasmic vestibule. All together our data indicate that the choline-bound cryo-EM structure of sfCHT reported here represents a substrate pre-release state.

**Figure 6.**
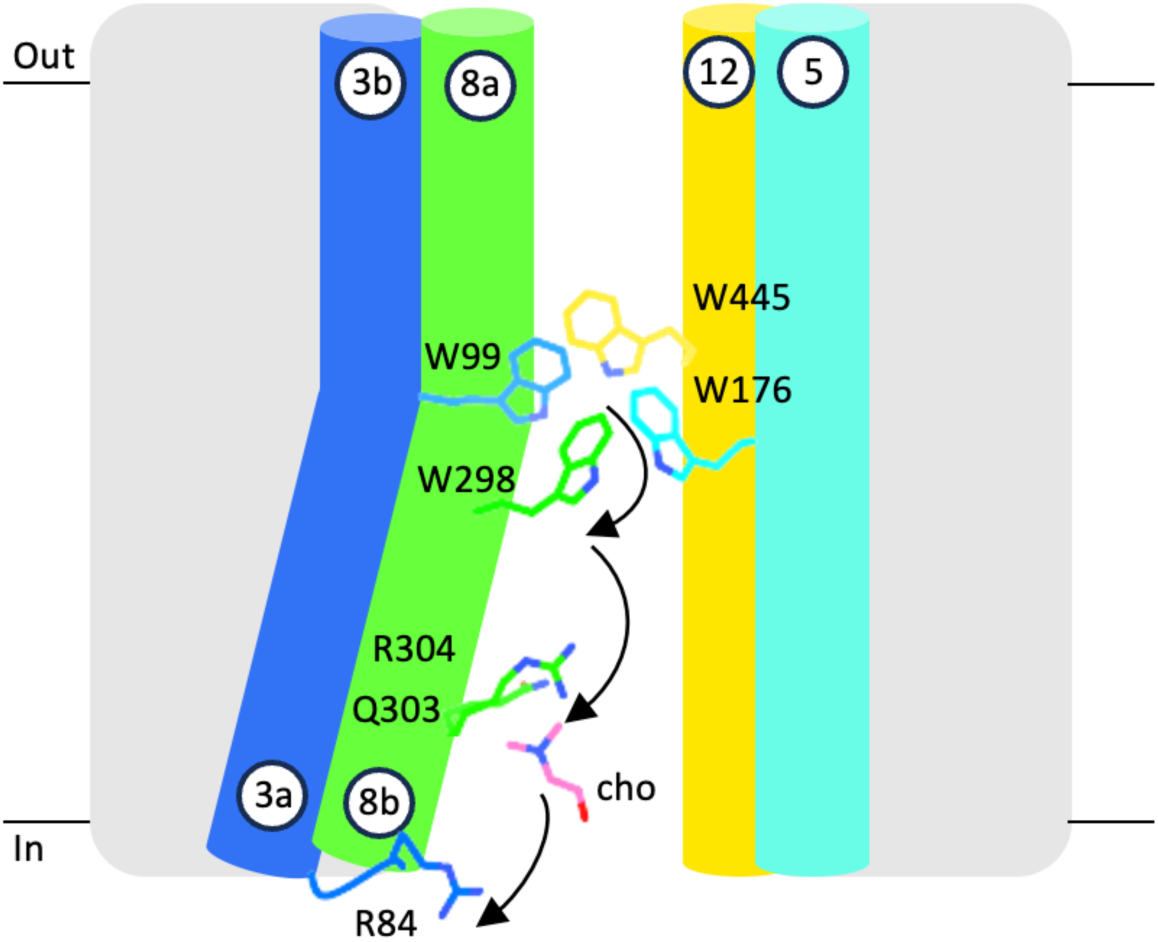
Model of choline translocation through CHT. Schematics of the inward-open cryo-EM structure of choline-bound sfCHT, with choline at the cytoplasmic site. Arrows indicate the stepwise movement of choline along the translocation pathway from the substrate-binding site to the cytoplasmic vestibule. Choline and the residues involved in local conformational changes that facilitate the movement of the substrate are depicted as sticks.

In our inward-facing structure in the presence of choline we could not find substrate in the putative substrate-binding-site between the unwound segments of TM helices 3 and 8, while the CHT1 structure captured the transporter in an occluded conformation with choline bound at the canonical substrate-binding site. We cannot discard that the detergent used for the purification of the protein stabilizes the transporter in an inward-facing conformation with a constricted substrate-binding site, thereby favoring choline binding at the cytoplasmic site. Despite the absence of choline in the central cavity of sfCHT, PELE analysis identified a minimal energy pose consistent with the choline-binding site in CHT1^9–11^ (**Supplementary Fig. 6b**). However, the absence of a Cl^-^ ion, which stabilizes choline by interacting with aromatic residues, may account for differences in binding pose. Whether Cl⁻ dependence emerged during evolution remains to be determined.

Although recent structural evidence has established the framework of substrate recognition mechanism of CHT1^9–11^, the translocation mechanism remains incompletely understood. Substitution analysis of CHT1 residues homologous to those implicated in sfCHT translocation reveals their critical role (**Fig. 5**). Substitution of residues within the substrate-binding site to alanine completely abolished transport in CHT1, highlighting their critical role in choline recognition and/or translocation. Interestingly, conservative substitutions to other bulky residues partially preserved transporter activity, emphasizing the importance of maintaining steric bulk within the substrate-binding site. These conserved aromatic residues might serve to confine the small choline molecule within a well-defined binding pocket, enabling the substrate to adopt productive poses, interacting at the same time with the unwound segments of TM helices 3 and 8, key for efficient translocation^23,24^. A similar mechanism has been reported for other transporters of small substrates, such as GlyT1^29^ and Asc1^22^, where aromatic residues in the substrate-binding site play a crucial role in efficient substrate recognition. Additionally, our results suggest *W62* and *W141*, would function as intracellular gates adjacent to the substrate-binding site, while *W406* might serve as extracellular gate (**Fig. 5**). Substitution of residues in the intracellular tunnel severely impaired choline transport activity in CHT1, affecting *V_max_* but not *K_m_*, reinforcing the role of these residues in choline translocation. In particular, the bulky residues *R47* and *R260* were found to be essential for function. These results reveal the stepwise evolutionarily conserved mechanism for choline translocation through CHT1 (**Fig. 6, Supplementary movie 1**).

In summary, our investigations into the Na^+^-dependent choline transport of sfCHT and the comparative analysis with CHT1 has unveiled that the choline translocation mechanism is evolutionarily conserved in CHT between bacteria and human orthologs. Since bacteria such as *Salimicrobium flavidum* precedes animals in evolution, sfCHT likely represents a more ancestral version of CHT than animal CHTs or human CHT1, suggesting that the foundations of Na^+^-dependent high-affinity choline transport mechanism were laid early in evolution.

## Data availability

The cryo-EM density maps of Na^+^-bound and choline-bound sfCHT have been deposited in the Electron Microscopy Data Bank (EMDB) under the accession codes EMD-54347 and EMD-54346, respectively. The corresponding atomic coordinates have been deposited in the Protein Data Bank (PDB) under PDB IDs 9RWT and 9RWS. Source data are provided with this paper.

## Acknowledgements

Authors thank Professor Randy Blakely for kindly providing the plasmid encoding CHT1 variant LL/VV. Authors thank Professor Israel S. Fernández for assistance in cryo-specimen preparation. This work has been supported by grants PID2023-146771NA-I00 to I.T., PID2019-104423GB-I00 and PID2022-143177NB-I00 to I. U.-B, PID2022-137310NB-I00 to E. E-M. and PID2022-140351OB-I00 to V. G., all funded by Spanish Ministry of Science, Innovation and Universities, Spanish Research Agency and European Regional Development Fund (MICIU / AEI /10.13039/501100011033 / FEDER, UE). J. V-G. acknowledges financial support from a FPU2020 contract (ref. FPU20/02480) granted by the Spanish Ministry of Science and Innovation. H.J. acknowledges financial support received by the UPV/EHU under the call for the recruitment of predoctoral research staff in the UPV/EHU 2024, PIF24/133. JP. L-A acknowledges financial support from a PTA contract granted by the MCIN/AEI. J. P-L. acknowledges financial support from a predoctoral contract PREP2023-000235 granted by the Spanish Ministry of Science, Innovation and Universities, Spanish Research Agency and European Social Fund plus (MICIU / AEI /10.13039/501100011033 / FSE+). High-resolution cryo-EM data collection was performed at the Basque Resource for Electron Microscopy supported primarily by the Department of Education and the Innovation Fund of the Basque Government, the Fundación Biofísica Bizkaia, and the Spanish Ministry of Science and Innovation, through the Plan de Recuperación, Transformación y Resiliencia (PRTR) funded by the NextGenerationEU (PRTR-C17.I1) program.

## Author contributions

I.T. conceived the project. I.T., J. V-G., E. E-M., and I. U-B. designed research and planned experiments. I.T. and J. V-G. performed the bacterial homolog selection. J. V-G., H.J., J. P-L. and B. O-L. performed protein expression and purification. J. V-G. and H.J. performed cryo-EM specimen preparation. J. V-G., JP. L-A., H.J. and I.T. carried out cryo-EM study and built and refined the atomic models. J. V-G., I.T. and E. E-M. analysed and interpreted the structures. V.G. conducted PELE analysis. A. M-J., P.B. and E. E-M. performed the mutational and functional studies. J. V-G. and I.T. wrote the original draft of the manuscript. I.T., J. V-G., E. E-M., and I. U-B. edited the manuscript with additional input from all authors. All authors participated in discussions and provided ideas for the work.

## Supplementary information

Supplementary information includes Supplementary Figures 1-13 and Supplementary Movie 1.

## Materials & Correspondence

Correspondence and request of materials should be addressed to Igor Tascón, Iban Ubarretxena-Belandia: or Ekaitz Errasti-Murugarren

## Methods

### Sequence alignment

The sequence conservation of SLC5A7 was assessed using a list composed of *Homo sapiens* and ten bacterial homologous sequences obtained from the BLAST-NCBI server. *Salimicrobium flavidum*, *Pseudobacteriovorax antillogorgiicola*, *Cyclobacterium halophilum*, *Halopolyspora algeriensis*, *Algoriphagus chordae*, *Algoriphagus aquimarinus*, *Bacillus aryabhattai*, *Halobacillus aidingensis*, *Melghirimyces thermohalophilus*, and *Marininema mesophilum*. PROMALS3D^30^ was employed for multiple sequence alignment, enabling alignment based on both sequence and secondary structure prediction. Jalview^31^ was used for figure preparation.

### Homologs selection

The selected bacterial sequences were codon optimized for expression, synthesized and cloned into two types of pB24 expression vector by GenScript company, adding an N-terminal or a C-terminal His_10_ tag to the expressed protein, respectively, and transformed into *E. coli* BL21 cells. Cells were grown at 37 °C in 5 mL LB medium supplemented with 100 μg/mL ampicillin, and overexpression was induced at different OD_600_ by the addition of different amounts of arabinose ranging from 0.001% to 0.02% at 37 °C or 20 °C, and at different induction times. Afterwards, cells were harvested and resuspended in 20 mM Tris pH 7.4, 150 mM NaCl buffer, supplemented with 0.5 mM PMSF, 1 mM benzamidine, and DNase I. Cells were disrupted using sonication, and all subsequent steps were performed at 4 °C. Cell debris and unbroken cells were pelleted at 15,000 x g for 15 min. Then, the membrane pellet was resuspended in 20 mM Tris HCl pH 7.4, 150 mM NaCl buffer to a concentration of 100 mg/mL and solubilized by the addition of 1% DDM for 1 h. Afterwards, the solution was centrifuged at 180,000 x g for 30 min to pellet unsolubilized particles. The supernatant was incubated with Ni-NTA beads for 1 h in 20 mM Tris pH 7.4, 150 mM NaCl, 0.04% DDM, 10 mM imidazole buffer. Unbound and nonspecifically bound proteins were eluted with 20 mM Tris pH 7.4, 150 mM NaCl, 0.04% DDM, 50 mM imidazole. The protein was further purified by size exclusion chromatography using a Superdex200 Increase 10/300 GL column (GE Healthcare) previously equilibrated with 20 mM Tris pH 7.4, 150 mM NaCl, 0.02% DDM. The expression tests were evaluated with SDS-PAGE and nLC-MS/MS, and the best behaving targets were used for negative staining screening.

### Expression and purification of sfCHT

The gene encoding WT sfCHT was cloned into a pB24 expression vector, adding a C-terminal His_10_ tag to the expressed protein, and transformed into *E. coli* BL21 cells. An overnight preculture was grown at 37 °C in LB medium supplemented with 100 μg/mL ampicillin. This culture was diluted 100-fold in 4 L of fresh LB medium with 100 μg/mL ampicillin at 37 °C, and overexpression was induced by the addition of 0.01% arabinose when an OD_600_ of 1 was reached. After 2.5 h of induction at 37 °C, cells were harvested and resuspended in 20 mM Tris pH 7.4, 150 mM NaCl buffer, supplemented with 0.5 mM PMSF, 1 mM benzamidine, and DNase I. Cells were disrupted using an Emulsiflex-C5 high pressure homogenizer (Avestin), and all subsequent steps were performed at 4 °C. Cell debris and unbroken cells were pelleted at 15,000 x g for 15 min. Membranes were collected by centrifugation at 180,000 x g for 3 h. The membrane pellet was resuspended in 20 mM Tris pH 7.4, 150 mM NaCl buffer to a concentration of 100 mg/mL and solubilized by the addition of 1% DDM for 1 h. Afterwards, the solution was centrifuged at 180,000 x g for 30 min to pellet unsolubilized particles. The supernatant was incubated with Ni-NTA beads for 1 h in 20 mM Tris pH 7.4, 150 mM NaCl, 0.04% DDM, 10 mM imidazole buffer. Unbound and nonspecifically bound proteins were eluted with 20 mM Tris pH 7.4, 150 mM NaCl, 0.04% DDM, 50 mM imidazole. sfCHT was eluted by incubating the sfCHT-bound Ni-NTA beads with the protease 3C. The protein was further purified by size exclusion chromatography using a Superdex200 Increase 10/300 GL column (GE Healthcare) previously equilibrated with 50 mM Tris pH 7.4, 100 mM NaCl, 0.02% DDM. Fractions containing the protein sfCHT were pooled and concentrated for Cryo-EM sample preparation. The purification process was assessed using SDS-PAGE and nLC-MS/MS.

### Negative staining

During the screening of homologous proteins of CHT1, those targets that could be purified were screened in negative staining for its suitability for cryo-EM: good particle distribution, without protein aggregates, and sufficient protein concentration. Specimens of sfCHT at different concentrations in 50 mM Tris pH 7.4, 100 mM NaCl, 0.02% DDM buffer were incubated onto previously glow-discharged continuous carbon film 400-mesh copper grids for 2 min, then washed in buffer for 30 s and subsequently stained for 1 min with 2% (w/v) uranyl acetate at pH 5, previously filtered. The grids were blotted onto filter paper and loaded in a JEM 1400 Plus transmission electron microscope (JEOL Japan) operating at 100 kV with a sCMOS camera for image acquisition.

### Cryo-EM specimen preparation and data collection

Purified sfCHT was concentrated up to 4 mg/mL for cryo-EM sample preparation. In case of the choline bound structure, right before vitrification, choline was added to a final concentration of 1 mM. Vitrification was carried out in a ThermoFisher Vitrobot Mark IV double-side blotting automated plunge freezer at 4 °C and 100% humidity. 3 μL of the sample was applied onto a previously glow-discharged UltrAuFoil^®^ R 0.6/1 gold foil on gold 300 mesh grid. The grid was double-side blotted for 3.5-5 s, using blot force 0, and was plunge-frozen in liquid ethane. The specimen was imaged on a 300 kV Krios G4 (ThermoScientific), equipped with a BioContinuum/K3 camera (Gatan) operating in counting mode at a calibrated pixel size of 0.8238 Å/pix for Na^+^-bound sfCHT and 0.6462 Å/pix for choline-bound sfCHT. A defocus range between -1.0 μm and -1.6 μm was used, and movies were recorded with a maximum total accumulated exposure of 60 e^-^/Å^2^ fractionated into 60 frames in case of Na^+^-bound sfCHT or 50 e^-^/Å^2^ fractionated into 50 frames for choline-bound sfCHT. Movies were recorded automatically using EPU 2 (ThermoScientific) with Aberration Free Image Shift (AFIS) and Fringe-free imaging (FFI).

### Image processing

Initial frame alignment of the recorded movies and contrast transfer function (CTF) estimation of the micrographs was carried out with cryoSPARC live^32^. 18,603 and 21,999 micrographs were collected for Na^+^- and choline-bound sfCHT, respectively. Only micrographs with maximum resolution better than 3.8 Å according to CTF estimation were selected for further processing. Automatic picking in cryoSPARC live was performed firstly without references by searching for Gaussian signals and then 2D classification helped to select good 2D class averages as templates for picking particles with reference. Subsequent data processing was carried out using cryoSPARC. This initial dataset contained 6,671,231 and 5,260,773 particles for Na^+^- and choline-bound sfCHT, respectively, both extracted to bin 2. Afterwards, a harsh cleaning of particles was performed in 2D classification so only 2D class averages clearly showing TM helices were kept for further steps. These particles were used to build ab-inito models with a starting resolution of 15 Å, a maximum resolution of 7 Å, 300 images per minibatch at the beginning and 1000 images at the end. We considered a good initial reference those volumes that keep cryo-EM density for transmembrane helices after the density from the detergent micelle region fades out. Particles from discarded 2D class averages were used to build decoy models acting as initial references to trap bad particles to further clean the particle dataset during heterogeneous refinement. 3,640,162 and 3,297,559 particles corresponding to Na^+^- and choline-bound sfCHT, respectively, were used together with the good ab-initio model and three decoy models for two rounds of heterogeneous refinement, the first with an initial resolution of 10 Å and the second 8 Å. Afterwards, a non-uniform refinement was carried out, starting with an initial resolution of 15 Å. To improve resolution, the selected particles so far were employed as seeds for Topaz training and extraction, and a similar protocol was followed with the extracted particles. After merging the previously curated particle set with those selected using Topaz^33^ and removing duplicate particles, two different approaches were employed for Na^+^- and choline-bound sfCHT.

For Na^+^-bound sfCHT, an initial non-uniform refinement was performed with a starting resolution of 15 Å, followed by particle extraction to bin 1. A second round of non-uniform refinement was carried out, maintaining the initial resolution of 15 Å. This was followed by a local refinement with an initial resolution of 9 Å, using a mask for transmembrane helices. A 3D classification was then performed with an initial resolution of 15 Å and target resolution of 2.5 Å to further clean up the subset of particles. Finally, a non-uniform refinement was applied to the selected volume, using an initial resolution of 9 Å, a mask for the transmembrane helices, and minimizing over per-particle scale at each iteration of refinement.

In the case of choline-bound sfCHT, two rounds of heterogeneous, non-uniform and local refinement were conducted. The heterogeneous and non-uniform refinements were performed with an initial resolution of 15 Å, whereas the local refinement was carried out with an initial resolution of 9 Å and a mask focused on the transmembrane helices. Subsequently, particles were extracted to bin 1, and non-uniform and local refinements were performed again with the same parameters, incorporating the minimization over per-particle scale at each iteration of refinement.

This workflow was optimized after extensive testing to address the challenges of this dataset, which is particularly difficult due to the small particle size and the predominance of transmembrane helices within the micelle. While we tested common cryo-EM processing tools such as CTF refinement and reference-based motion correction, these did not improve the reconstruction. The described approach proved to be the most effective for this sample.

### Model building

In the case of model building of Na^+^-bound sfCHT, we generated an initial model using ModelAngelo^34^. After rigid-body fitting into our 2.83 Å resolution cryo-EM map in UCSF ChimeraX^35^ a Na^+^ ion was manually added using Coot^36^. Residues from 1 to 21, from 70 to 80, and the C-terminal 7 residues could not be assigned to any Cryo-EM density, so were removed from the model. A final real-space refinement in Phenix was performed to improve the fit and to optimize stereochemistry^37^. For model building of choline-bound sfCHT, the Na^+^-bound sfCHT model was rigid-body fitted into our 3.28 Å resolution cryo-EM map in UCSF ChimeraX^35^. Na^+^, H_2_O and choline molecules were subsequently added using Coot^36^. Following this, a final real-space refinement was carried out using Phenix to obtain an optimal geometry model^37^.

### Reconstitution into proteoliposomes (PLs)

WT and variant sfCHT were reconstituted in *E. coli* polar lipids (Avanti Lipids), as previously described^38^. In brief, lipid was dried under N_2_ and suspended in reconstitution buffer containing 20 mM Tris pH 7.4, 150 mM KCl. The suspension was then sonicated to clarity, and purified sfCHT was added to a 1:50 protein/lipid ratio (w/w). Then, liposomes were destabilized by the addition of 1.25% β-D-octylglucoside (OG) and incubated in ice with occasional agitation for 5 min. DDM and OG were removed by dialysis for 40 h at 4°C against 100 volumes of dialysis buffer (20 mM Tris pH 7.4, 150 mM KCl). PL suspensions were frozen in liquid N_2_ and stored at -80°C until use.

### Choline transport assays in PLs

Choline uptake assays were initiated after mixing 20 μl of cold PLs with 180 μl of sodium or sodium-free transport buffer (10 mM HEPES pH 7.4, 137 mM NaCl or 137 mM KCl, 5 mM KCl, 2 mM CaCl_2_, 1 mM MgSO_4_), 1 μCi/mL radiolabelled choline chloride (Perkin Elmer, Waltham, USA), and unlabelled choline chloride to the desired concentration. Transport experiments were performed at room temperature for the indicated periods of time and were stopped by the addition of 2 mL of ice-cold stop buffer (5 mM choline chloride in transport buffer) and filtration through 0.45 μm pore size membrane filters (Sartorius Stedim Biotech). Filters were then washed two times with 2 mL of cold stop buffer and dried, and the trapped radioactivity was counted. Transport measurements are normalized to the sfCHT protein concentration measured for each sfCHT-PL batch. Data are expressed as the mean ± S.D. of three experiments performed on different days and on different reconstitutions.

### PELE

PELE (Protein Energy Landscape Exploration) is a Monte Carlo (MC) molecular modeling software capable of mapping complex intermolecular biophysical problems, such as ligand migration, binding site search, local induced fit, etc.^39^. At each MC step PELE performs multiple sampling routines including: i) ligand random translation and rotation, ii) protein backbone motion along a randomly chosen normal mode, iii) side chain sampling of all amino acids within 6 Å from the ligand, and iv) and overall minimization of the whole system. PELE global exploration^40^ (also known as SiteFinder) was first used as an unsupervised docking of choline from outside of the transporter to explore different poses. The protocol involved 255 computing cores for 24 h, where each core performed a (search) trajectory that was randomly placed at the surface of the transporter. We found a minimum energy pose at the intracellular tunnel exit (**Fig. 3b**). From this position, a series of successive local induced fit simulations were run to identify potential inner binding poses along the intracellular tunnel and within the substrate-binding site. Each induced fit simulation was allowed to explore a 9 Å sphere around the initial ligand position, involving 128 computing cores for 6 h.

### Mutagenesis and transfection of CHT1 variants

HeLa cells were maintained at 37 °C in a humidified 5% CO_2_ environment in DMEM supplemented with 10% fetal bovine serum, 50 units/mL penicillin, 50 μg/mL streptomycin and 2 mM L-glutamine. HeLa cells were transiently transfected in a 24-well plate with 400 ng/well with the endocytosis resistant human CHT1-LV/AA variant^41^ (kindly provided by Professor Randy Blakely, Florida Atlantic University, USA) or CHT1 variants using Lipofectamine 2000 (Invitrogen, Carlsbad, USA). Single point substitutions were introduced using the QuikChange mutagenesis kit (Stratagene, San Diego, USA). All substitutions were verified by sequencing. Choline transport assays were carried out 24 h after transfection. Endocytosis resistant human CHT1-LV/AA variant, we refer as WT CHT1 in the context of this study, was used as template for the generation of CHT1 variants.

### Visualization of CHT1 variants by fluorescence microscopy

To analyze the effect of the substitutions on CHT1 protein expression and plasma membrane localization, fluorescence microscopy of wild type and mutant transporters was performed on a semiconfluent monolayer of transfected HeLa cells cultured on glass coverslips. Glass coverslip-grown cells were rinsed three times with phosphate-buffered saline-Ca^2+^-Mg^2+^ and fixed for 5 min in ice-cold methanol. Fixed cells were blocked and permeabilized in blocking buffer (4% FBS and 0.5% Triton X-100 in PBS) for 1 h and then incubated for 1 h with primary antibody (anti-CHT, #CHT-Go-Af890, Nittobo Medical). Secondary donkey-anti-goat-Alexa 546 antibody (Life Technologies) was incubated for 2 h protected from light and rinsed three times with phosphate-buffered saline. Nuclear staining was performed by incubating 1 µg/mL Hoechst (Thermo Fisher Scientific) for 10 min, rinsed three times with phosphate-buffered saline and then mounted with aqua-poly/mount coverslipping medium (Polysciences Inc.). Images were taken using a Nikon E1000 upright epifluorescence microscope. All images were captured during 200 ms.

### Choline transport assays in HeLa cells

Choline uptake was measured in WT and variant CHT1, and mock-transfected HeLa cells, as previously described^22^, by exposing replicate cultures at room temperature to [^3^H]-labelled choline chloride (1 μCi/mL; Perkin Elmer, Waltham, USA) in transport buffer (10 mM HEPES pH 7.4, 137 mM NaCl, 5 mM KCl, 2 mM CaCl_2_, 1 mM MgSO_4_). Initial rates of transport were determined using an incubation period of 2 min, as described^41^. Assays were terminated by washing with an excess volume of ice-cold stop buffer (5 mM choline chloride in transport buffer). Transporter-mediated choline uptake was calculated by subtracting the uptake measured in mock-transfected cells.

For saturation kinetics, transfected cells were incubated with 1 μCi/mL [^3^H] choline chloride and varying concentrations of unlabelled choline chloride (0–100 or 100-500 μM, depending on the mutant to be analysed). Cold substrates were prepared at 100 mM, aliquoted and stored at -20 °C until use. Aliquots were thaw only once to reduce variability. Each replicate of the kinetic studies was performed simultaneously for WT and variant CHT1. The Michaelis-Menten and Eadie-Hofstee equations were then applied, and the kinetic parameters derived from this method were confirmed by linear regression analysis of the derived Eadie-Hofstee plots using the GraphPad Prism software. Data are expressed as the mean ± S.D. of three experiments performed on different days and on different batches of cells.

## References

1 Wargo, M. J. Homeostasis and catabolism of choline and glycine betaine: lessons from Pseudomonas aeruginosa. Appl. Environ. Microbiol. 79, 2112–2120 (2013). 10.1128/AEM.03565-12

2 Zeisel, S. H. Choline: essential for brain development and function. Adv Pediatr 44, 263–295 (1997).

3 Zeisel, S. H. Choline: critical role during fetal development and dietary requirements in adults. Annu. Rev. Nutr. 26, 229–250 (2006). 10.1146/annurev.nutr.26.061505.111156

4 Geiger, O., Lopez-Lara, I. M. & Sohlenkamp, C. Phosphatidylcholine biosynthesis and function in bacteria. Biochim Biophys Acta 1831, 503–513 (2013). 10.1016/j.bbalip.2012.08.009

5 Ojiakor, O. A. & Rylett, R. J. Modulation of sodium-coupled choline transporter CHT function in health and disease. Neurochem. Int. 140, 104810 (2020). 10.1016/j.neuint.2020.104810

6 Apparsundaram, S., Ferguson, S. M., George, A. L., Jr. & Blakely, R. D. Molecular cloning of a human, hemicholinium-3-sensitive choline transporter. Biochem. Biophys. Res. Commun. 276, 862–867 (2000). 10.1006/bbrc.2000.3561

7 Okuda, T. & Haga, T. Functional characterization of the human high-affinity choline transporter. FEBS Lett. 484, 92–97 (2000). 10.1016/s0014-5793(00)02134-7

8 Iwamoto, H., Blakely, R. D. & De Felice, L. J. Na+, Cl-, and pH dependence of the human choline transporter (hCHT) in Xenopus oocytes: the proton inactivation hypothesis of hCHT in synaptic vesicles. J. Neurosci. 26, 9851–9859 (2006). 10.1523/JNEUROSCI.1862-06.2006

9 Qiu, Y., Gao, Y., Huang, B., Bai, Q. & Zhao, Y. Transport mechanism of presynaptic high-affinity choline uptake by CHT1. Nat. Struct. Mol. Biol. 31, 701–709 (2024). 10.1038/s41594-024-01259-w

10 Xue, J., Chen, H., Wang, Y. & Jiang, Y. Structural mechanisms of human sodium-coupled high-affinity choline transporter CHT1. Cell Discov 10, 116 (2024). 10.1038/s41421-024-00731-7

11 Qiu, Y., Gao, Y., Bai, Q. & Zhao, Y. Ion coupling and inhibitory mechanisms of the human presynaptic high-affinity choline transporter CHT1. Structure (2024). 10.1016/j.str.2024.11.009

12 Ziegler, C., Bremer, E. & Kramer, R. The BCCT family of carriers: from physiology to crystal structure. Mol. Microbiol. 78, 13–34 (2010). 10.1111/j.1365-2958.2010.07332.x

13 Salvano, M. A., Lisa, T. A. & Domenech, C. E. Choline transport in Pseudomonas aeruginosa. Mol. Cell. Biochem. 85, 81–89 (1989). 10.1007/BF00223517

14 Wargo, M. J., Ho, T. C., Gross, M. J., Whittaker, L. A. & Hogan, D. A. GbdR regulates Pseudomonas aeruginosa plcH and pchP transcription in response to choline catabolites. Infect. Immun. 77, 1103–1111 (2009). 10.1128/IAI.01008-08

15 Oswald, C. et al. Crystal structures of the choline/acetylcholine substrate-binding protein ChoX from Sinorhizobium meliloti in the liganded and unliganded-closed states. J. Biol. Chem. 283, 32848–32859 (2008). 10.1074/jbc.M806021200

16 Perez, C. et al. Substrate specificity and ion coupling in the Na+/betaine symporter BetP. EMBO J. 30, 1221–1229 (2011). 10.1038/emboj.2011.46

17 Du, Y. et al. Structures of the substrate-binding protein provide insights into the multiple compatible solute binding specificities of the Bacillus subtilis ABC transporter OpuC. Biochem. J. 436, 283–289 (2011). 10.1042/BJ20102097

18 Pittelkow, M., Tschapek, B., Smits, S. H., Schmitt, L. & Bremer, E. The crystal structure of the substrate-binding protein OpuBC from Bacillus subtilis in complex with choline. J. Mol. Biol. 411, 53–67 (2011). 10.1016/j.jmb.2011.05.037

19 Barland, N. et al. Mechanistic basis of choline import involved in teichoic acids and lipopolysaccharide modification. Sci Adv 8, eabm1122 (2022). 10.1126/sciadv.abm1122

20 Yang, T. et al. Structure and mechanism of the osmoregulated choline transporter BetT. Sci Adv 10, eado6229 (2024). 10.1126/sciadv.ado6229

21 Shi, Y. Common folds and transport mechanisms of secondary active transporters. Annu Rev Biophys 42, 51–72 (2013). 10.1146/annurev-biophys-083012-130429

22 Rullo-Tubau, J. et al. Structure and mechanisms of transport of human Asc1/CD98hc amino acid transporter. Nat Commun 15, 2986 (2024). 10.1038/s41467-024-47385-3

23 Rodriguez, C. F. et al. Structural basis for substrate specificity of heteromeric transporters of neutral amino acids. Proc Natl Acad Sci U S A 118 (2021). 10.1073/pnas.2113573118

24 Errasti-Murugarren, E. et al. L amino acid transporter structure and molecular bases for the asymmetry of substrate interaction. Nat Commun 10, 1807 (2019). 10.1038/s41467-019-09837-z

25 Tascon, I. et al. Structural basis of proton-coupled potassium transport in the KUP family. Nat Commun 11, 626 (2020). 10.1038/s41467-020-14441-7

26 Weyand, S. et al. The alternating access mechanism of transport as observed in the sodium-hydantoin transporter Mhp1. J Synchrotron Radiat 18, 20–23 (2011). 10.1107/S0909049510032449

27 Jardetzky, O. Protein dynamics and conformational transitions in allosteric proteins. Prog Biophys Mol Biol 65, 171–219 (1996).

28 Hiraizumi, M. et al. Transport and inhibition mechanism of the human SGLT2-MAP17 glucose transporter. Nat. Struct. Mol. Biol. (2023). 10.1038/s41594-023-01134-0

29 Wei, Y. et al. Transport mechanism and pharmacology of the human GlyT1. Cell 187, 1719–1732 e1714 (2024). 10.1016/j.cell.2024.02.026

30 Pei, J., Kim, B. H. & Grishin, N. V. PROMALS3D: a tool for multiple protein sequence and structure alignments. Nucleic Acids Res. 36, 2295–2300 (2008). 10.1093/nar/gkn072

31 Waterhouse, A. M., Procter, J. B., Martin, D. M., Clamp, M. & Barton, G. J. Jalview Version 2--a multiple sequence alignment editor and analysis workbench. Bioinformatics 25, 1189-1191 (2009). 10.1093/bioinformatics/btp033

32 Punjani, A., Rubinstein, J. L., Fleet, D. J. & Brubaker, M. A. cryoSPARC: algorithms for rapid unsupervised cryo-EM structure determination. Nat. Meth. 14, 290–296 (2017). 10.1038/nmeth.4169

33 Bepler, T. et al. Positive-unlabeled convolutional neural networks for particle picking in cryo-electron micrographs. Nat. Methods 16, 1153–1160 (2019). 10.1038/s41592-019-0575-8

34 Jamali, K. et al. Automated model building and protein identification in cryo-EM maps. Nature (2024). 10.1038/s41586-024-07215-4

35 Goddard, T. D. et al. UCSF ChimeraX: Meeting modern challenges in visualization and analysis. Protein Sci. 27, 14–25 (2018). 10.1002/pro.3235

36 Emsley, P. & Cowtan, K. Coot: model-building tools for molecular graphics. Acta Crystallogr D Biol Crystallogr 60, 2126–2132 (2004). 10.1107/S0907444904019158

37 Adams, P. D. et al. PHENIX: a comprehensive Python-based system for macromolecular structure solution. Acta Crystallogr D Biol Crystallogr 66, 213–221 (2010). 10.1107/S0907444909052925

38 Kowalczyk, L. et al. Molecular basis of substrate-induced permeation by an amino acid antiporter. Proc Natl Acad Sci U S A 108, 3935–3940 (2011). 10.1073/pnas.1018081108

39 Lecina, D., Gilabert, J. F. & Guallar, V. Adaptive simulations, towards interactive protein-ligand modeling. Sci Rep 7, 8466 (2017). 10.1038/s41598-017-08445-5

40 Acebes, S. et al. Rational Enzyme Engineering Through Biophysical and Biochemical Modeling. Acs Catalysis 6, 1624–1629 (2016). 10.1021/acscatal.6b00028

41 Ruggiero, A. M. et al. Nonoisotopic assay for the presynaptic choline transporter reveals capacity for allosteric modulation of choline uptake. ACS Chem Neurosci 3, 767–781 (2012). 10.1021/cn3000718

