## Supplementary Information for "Structural insights into a conserved mechanism of choline translocation through CHT"

Jesus Vilchez-Garcia *et al.*

**This Supplementary Information includes:**

Supplementary Figures 1 to 13
Supplementary Movie S1

**Other Supplementary Materials for this manuscript include the following:**

Movie S1 (mp4 file)

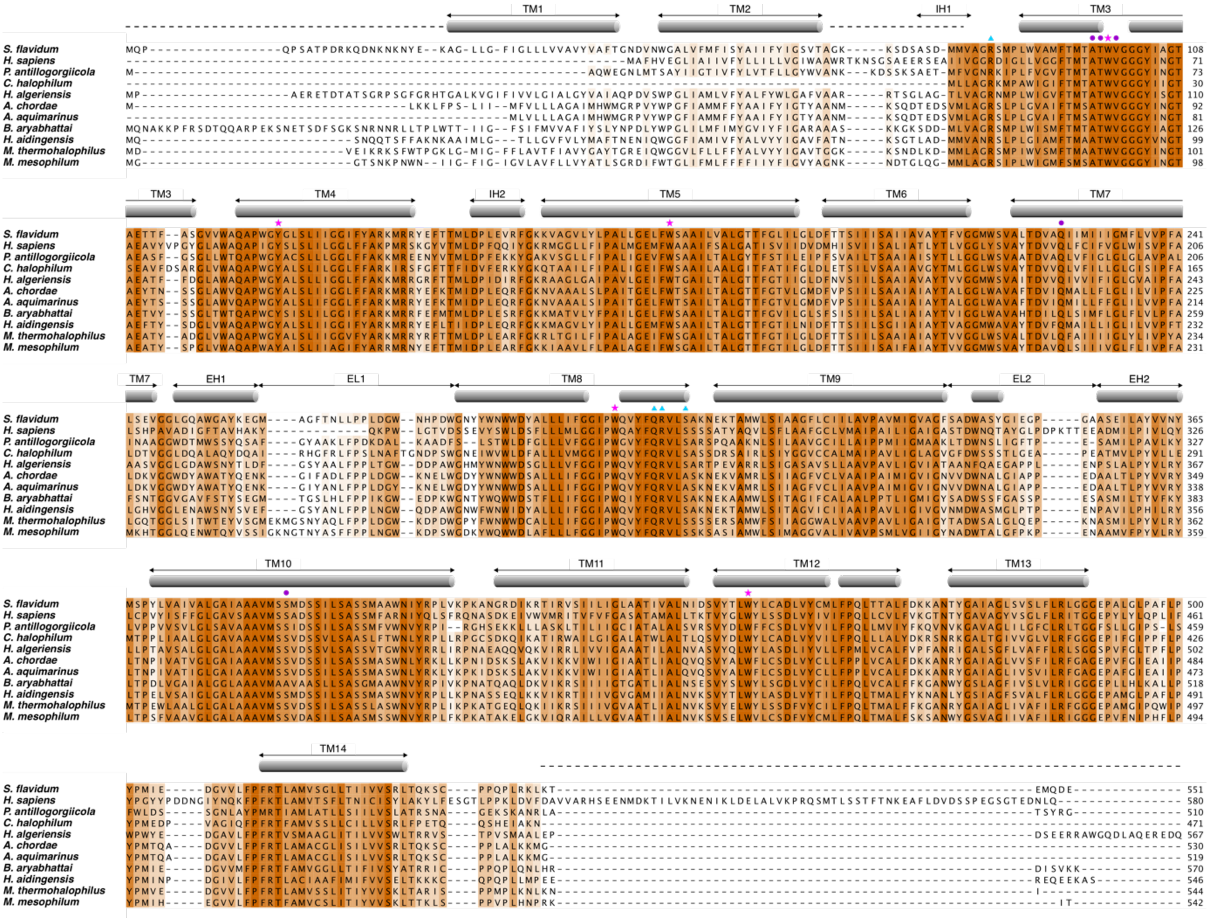


Supplementary Figure 1. **Sequence alignment of SLC5A7 from *S. flavidum* with human SLC5A7 and other homologous proteins from different bacteria.** Sequences of SLC5A7 from *Salimicrobium flavidum*, *Homo sapiens*, *Pseudobacteriovorax antillogorgiicola*, *Cyclobacterium halophilum*, *Halopolyspora algeriensis*, *Algoriphagus chordae*, *Algoriphagus aquimarinus*, *Bacillus aryabhattai*, *Halobacillus aidingensis*, *Melghirimyces thermohalophilus*, *Marininema mesophilum* are aligned. The sequence alignment was carried out using PROMALS3D, and the figure was prepared using Jalview. Sequence conservation is shown by coloring, with dark orange indicating full conservation. The secondary structure of sfCHT is shown with gray cylinders. Dashed lines before K22, between A69 and V81, and after L544 indicate unresolved residues in our structures due to protein flexibility in these segments. Magenta stars indicate residues constituting the substrate-binding site, purple circles residues involved in $Na^+$ coordination, and blue triangles indicate the identified key residues R84, Q303, R304 and S307. TM, IH, EH and EL stand for transmembrane helix, intracellular helix, extracellular helix and extracellular loop, respectively.

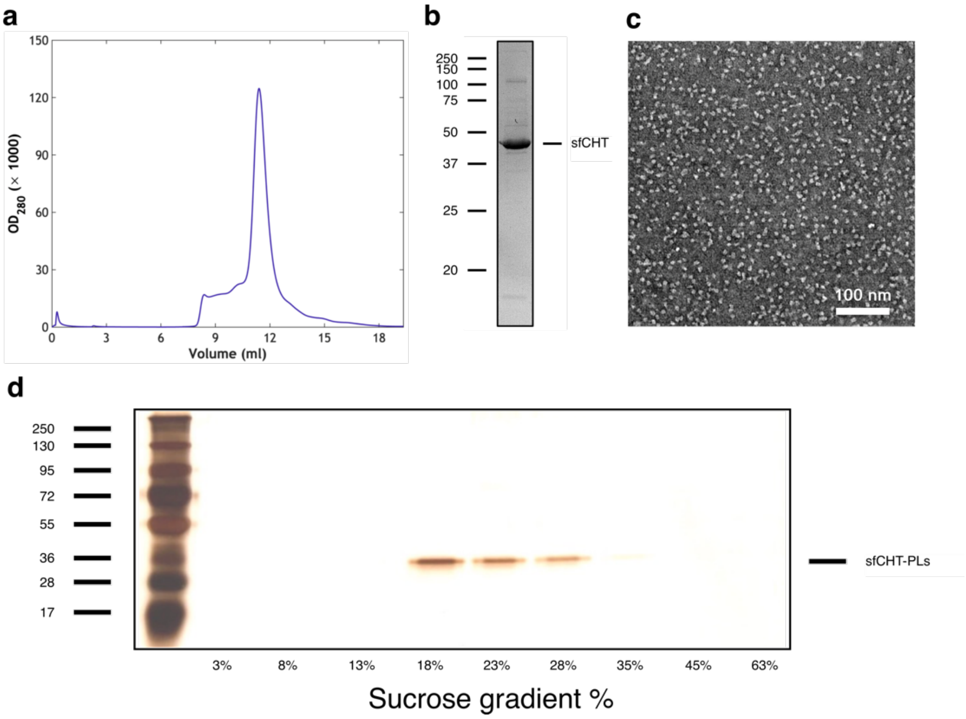


Supplementary Figure 2. **sfCHT for structural biology studies. a)** Size exclusion chromatography profile of a representative purification of sfCHT. **b)** SDS-PAGE showing a prominent band corresponding to purified sfCHT after size exclusion chromatography. **c)** Representative negative staining micrograph of sfCHT illustrating a good particle distribution without protein aggregates. **d)** Sucrose gradient of DDM-purified sfCHT and reconstituted in *E. coli* polar lipid liposomes. The layers of the gradient were analyzed by SDS-PAGE and further silver staining. Given concentrations correspond to sucrose concentrations before centrifugation.

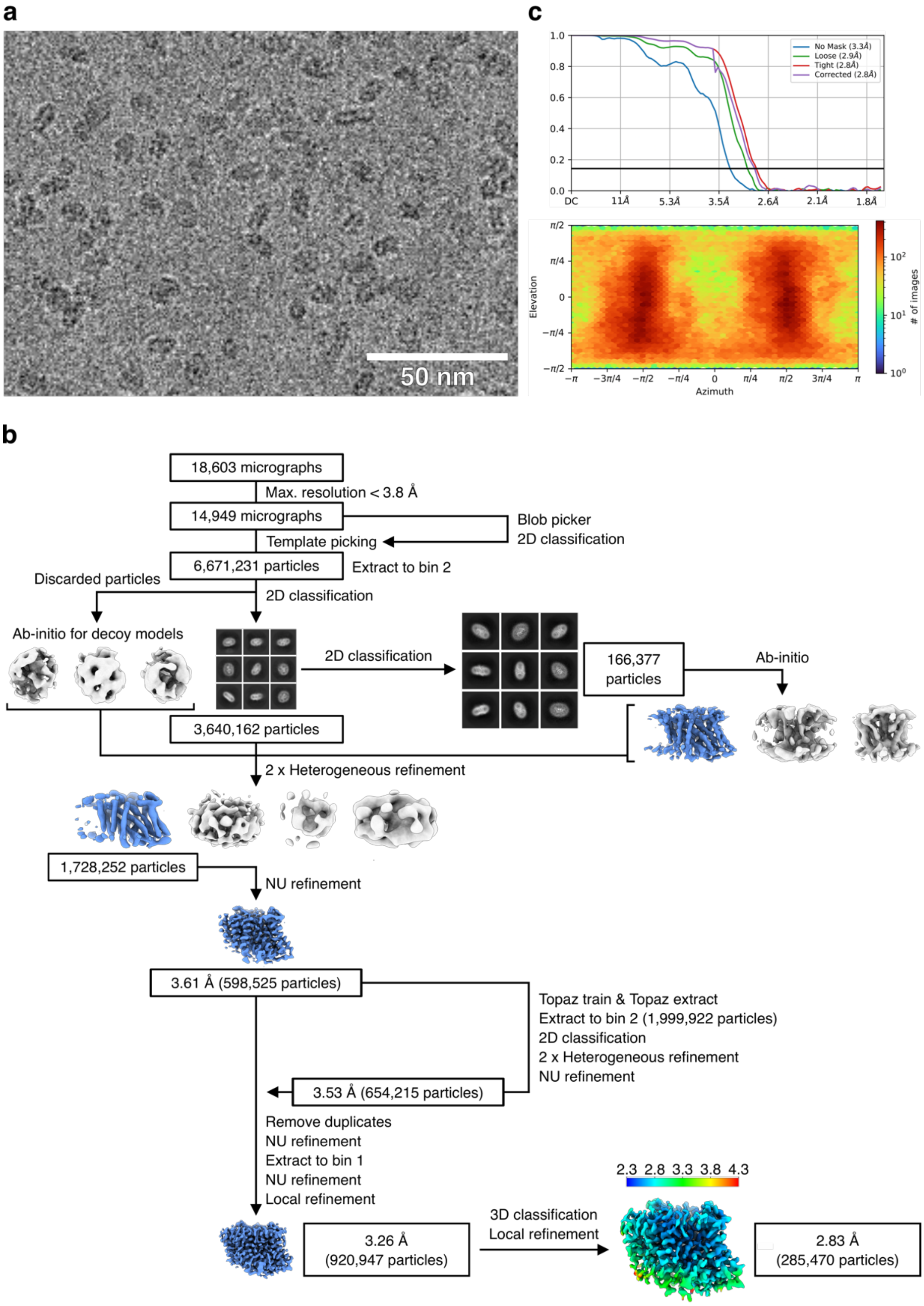


Supplementary Figure 3. **Cryo-EM data processing pipeline for Na$^+$-bound sfCHT. a)** Zoomed-in view of a representative cryo-EM micrograph of Na$^+$-bound sfCHT. **b)** Cryo-EM data processing workflow. **c)** Gold-standard FSC curve (top) and Euler diagram showing particle orientation distribution (bottom).

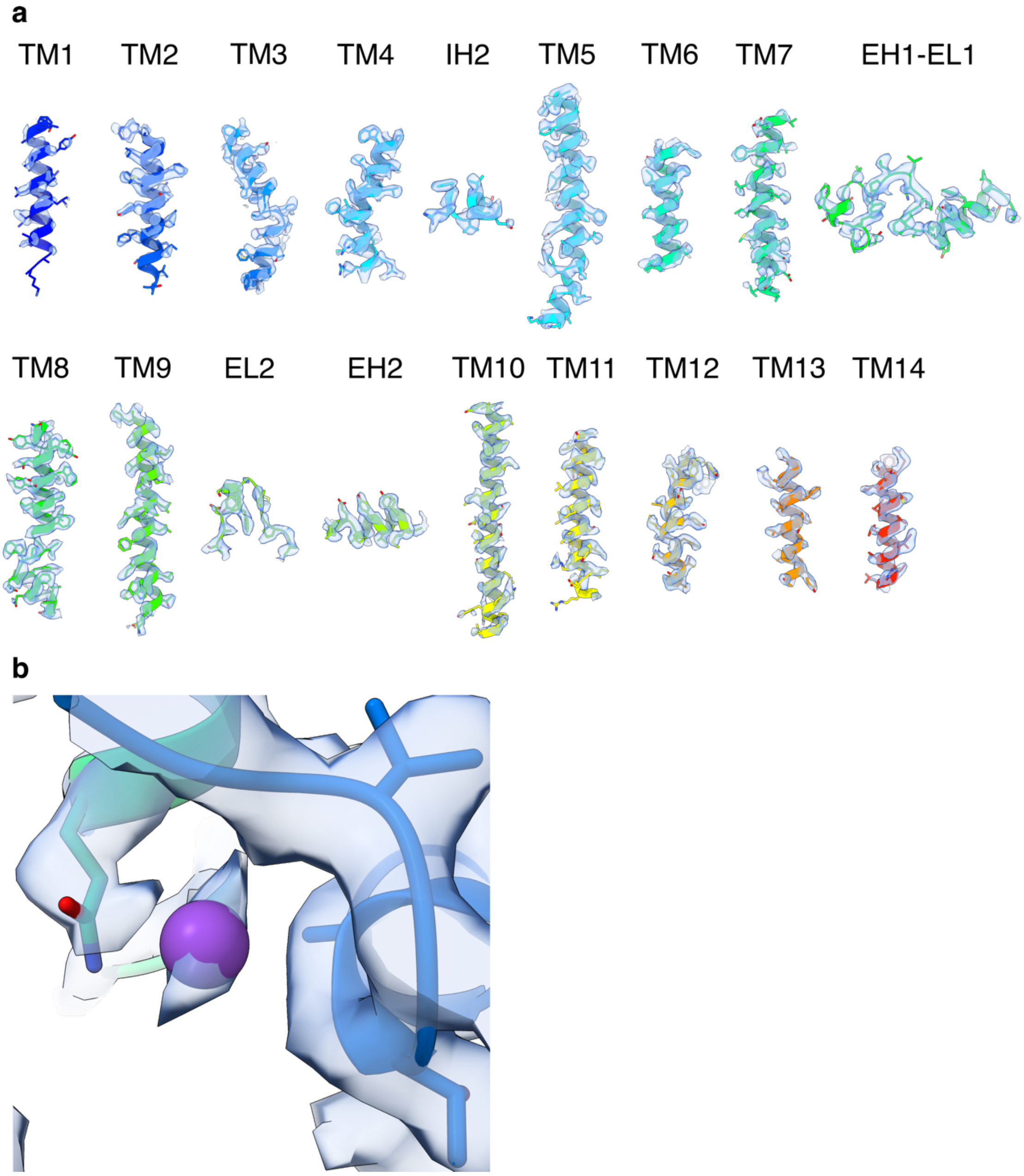


Supplementary Figure 4. **Cryo-EM map and model-to-map fit of $Na^+$-bound sfCHT.** The map threshold was set to an RMSD of 8.20 Å in all panels. **a)** The different segments of sfCHT, named as in Supplementary Figure 1, are depicted in cartoon format with the side chains shown in sticks. The cryo-EM density map is displayed around these segments. **b)** Close-up view of the $Na^+$ cation shown as spheres, interacting with residues depicted in sticks.

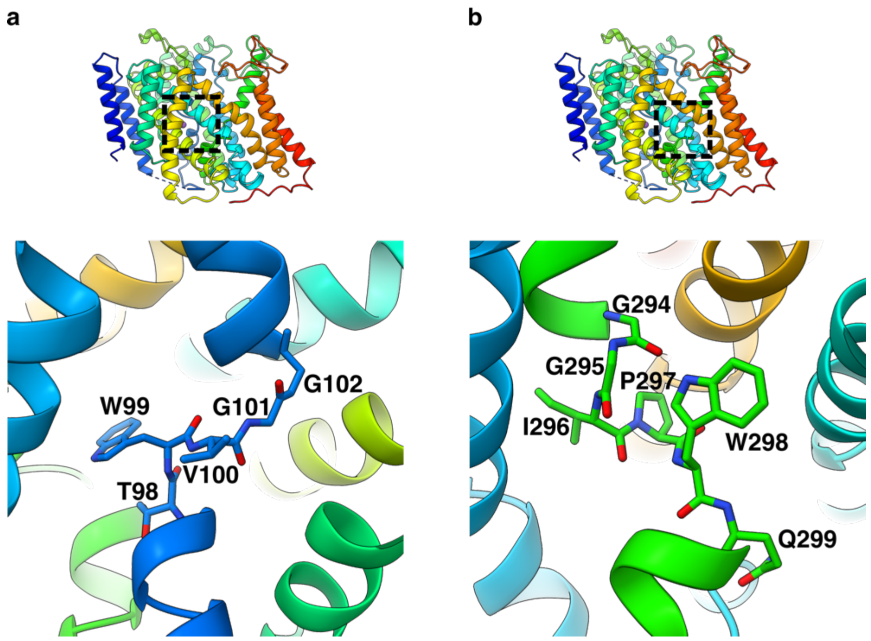


Supplementary Figure 5. **Discontinuities in TM helix 3 (a) and TM helix 8 (b)**. The unwound segments are depicted as sticks. The upper section of each panel highlights the specific region of the protein shown in the enlarged view below.

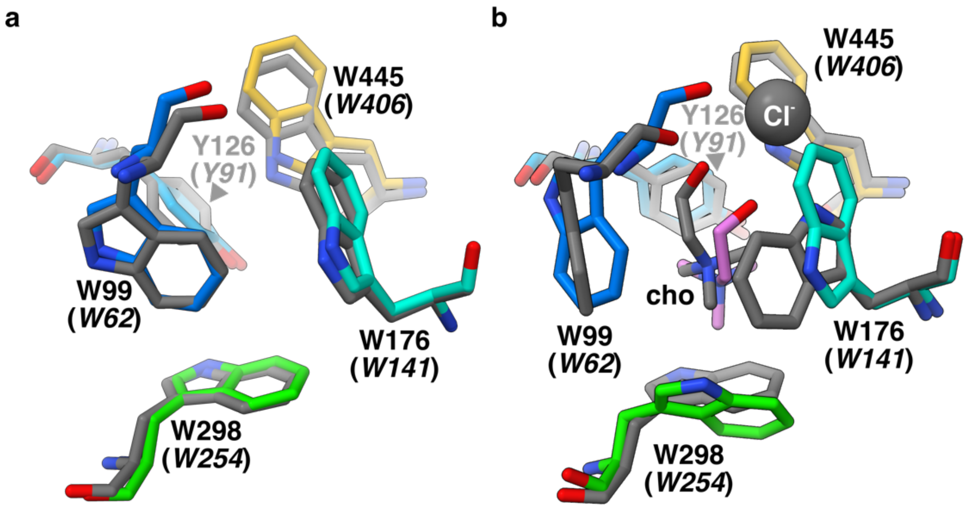


Supplementary Figure 6. **Substrate-binding site comparison between sfCHT and CHT1. a)** Structural alignment of inward-facing $Na^+$-bound sfCHT (colored) and inward-open apo CHT1 (PDB: 9BFI, shown in gray). **b)** Structural alignment of the PELE-predicted model of sfCHT with choline-bound to the substrate-binding site (colored) and the inward-facing CHT1 structure bound to $Na^+$, $Cl^-$ and choline (PDB: 9BFJ, shown in gray).

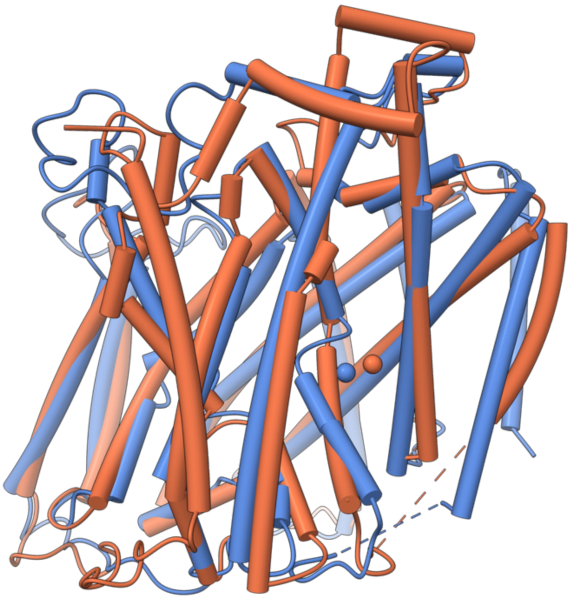

Supplementary Figure 7. **Structural homology between sfCHT and vSGLT.** Side view of superimposed sfCHT (in blue) and vSGLT (in coral), with α-helices displayed as tubes, and the $Na^+$ ions as spheres.

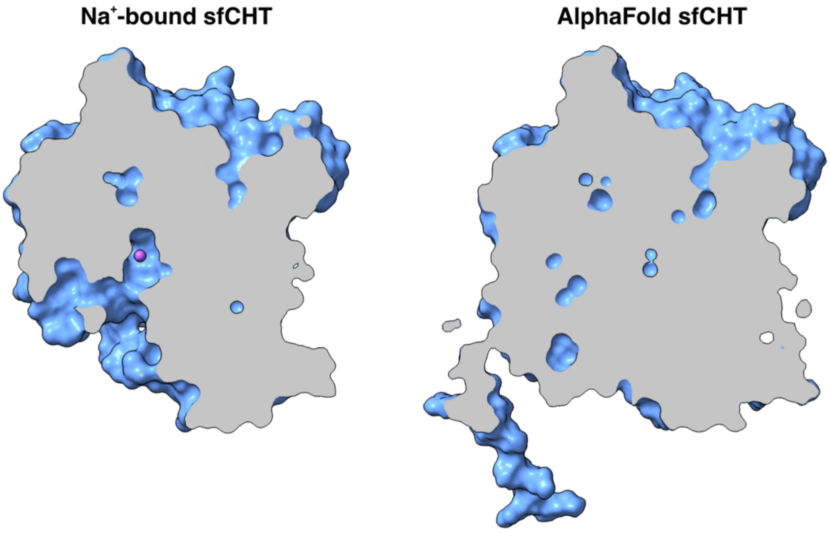


Supplementary Figure 8. **Accessibility of the substrate-binding site from the cytoplasm in $Na^+$-bound and AlphaFold sfCHT structures.** The panels illustrate a cross-section of each structure, revealing the internal architecture from a front view. The structures are represented as surface models. The $Na^+$ ion is shown as a purple sphere. In the inward-facing $Na^+$-bound sfCHT structure a tunnel from the cation to the inner side of the membrane is observed, whereas the AlphaFold sfCHT structure lacks any discernible tunnel.

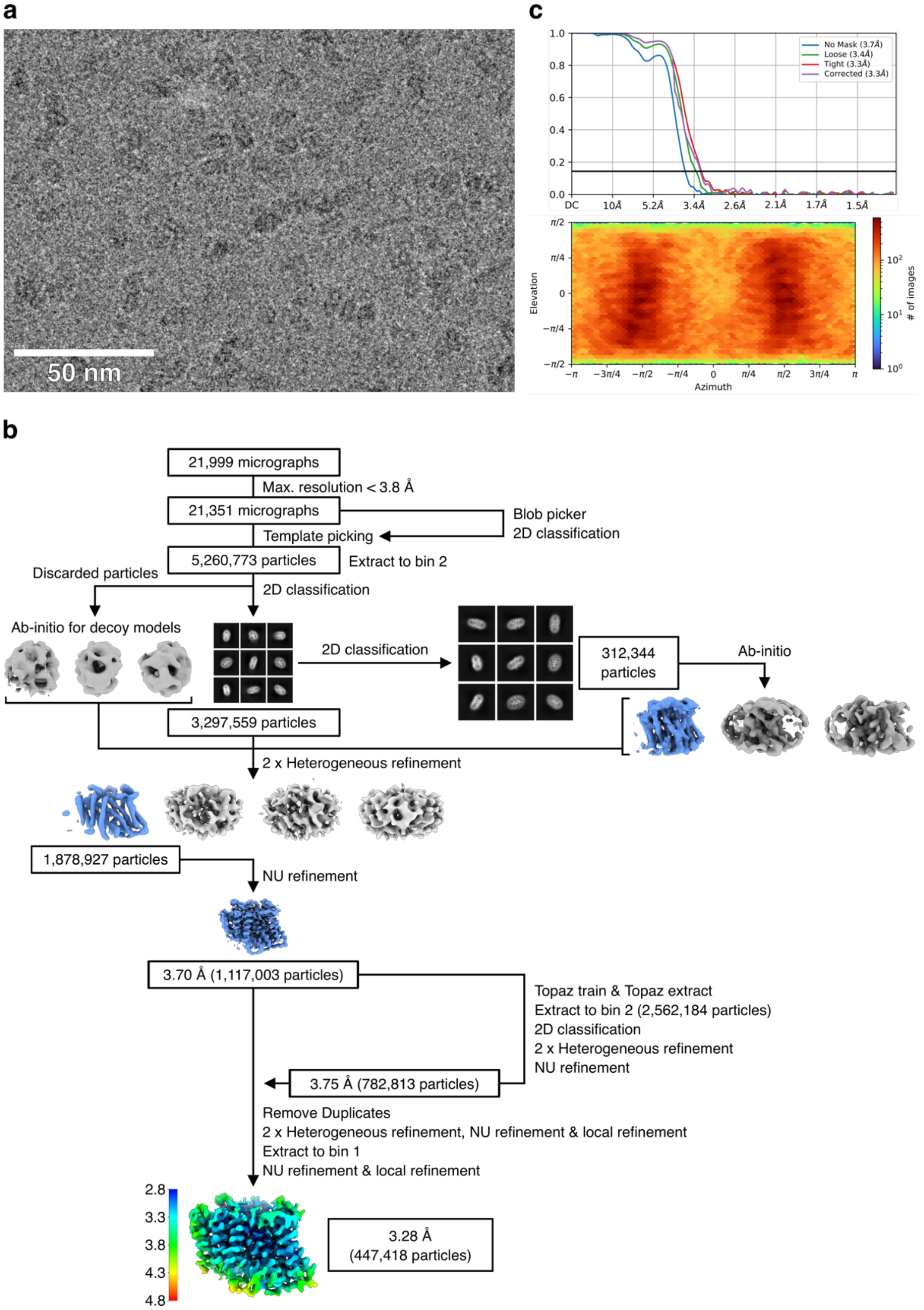


Supplementary Figure 9. **Cryo-EM data processing pipeline for choline-bound sfCHT. a)** Zoomed-in view of a representative cryo-EM micrograph of choline-bound sfCHT. **b)** Cryo-EM data processing workflow. **c)** Gold-standard FSC curve (top) and Euler diagram showing particle orientation distribution (bottom).

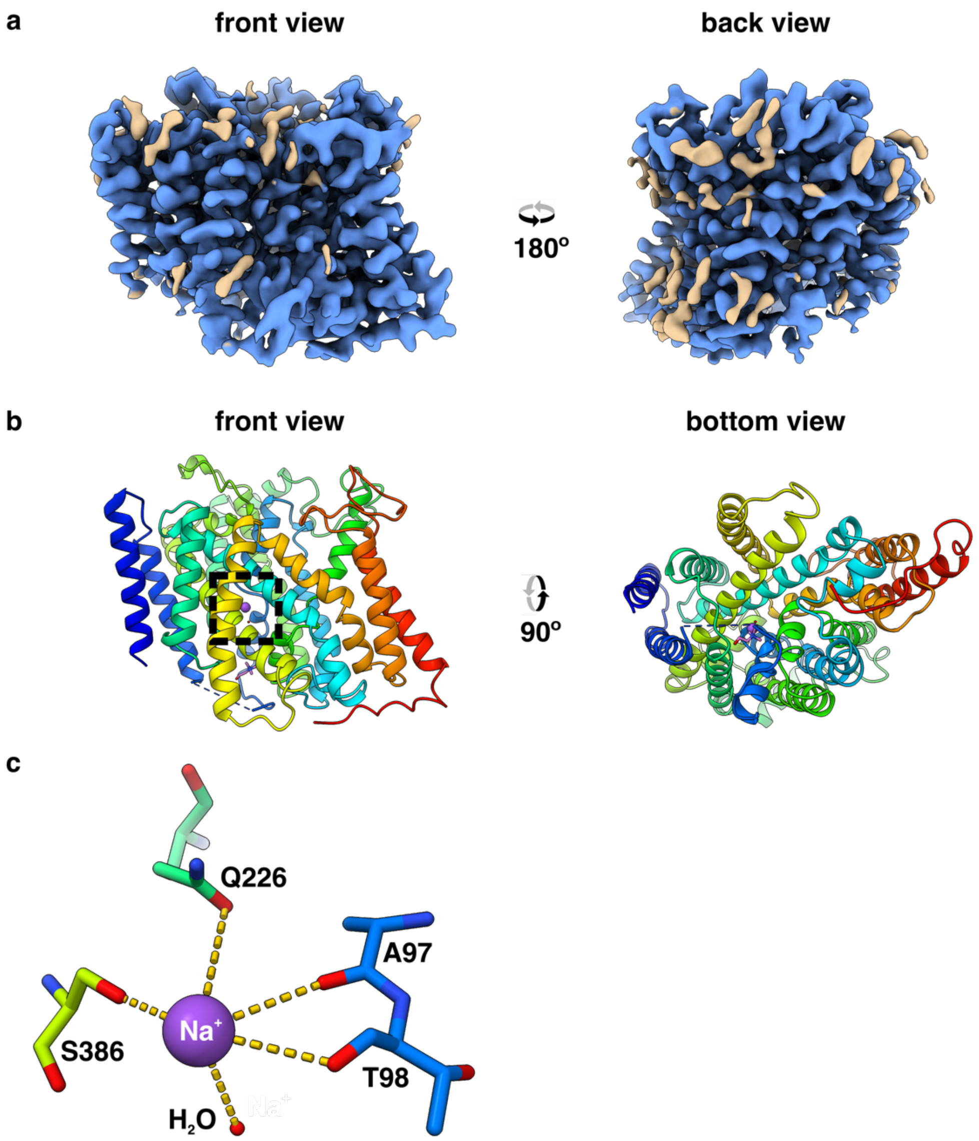


Supplementary Figure 10. **Overview of the structure of choline-bound sfCHT. a)** Cryo-EM map of sfCHT at 3.28 Å nominal resolution viewed from the front and back side. The map regions colored in blue correspond to protein and ligand-assigned densities, whereas non-protein unassigned densities are brown colored. **b)** Ribbon representation of the atomic model of sfCHT viewed from the membrane, colored in a rainbow gradient from blue (N-terminus) to red (C-terminus). The missing loop between TM helices 2 and 3 is depicted as a dashed line. $Na^+$ cation is shown as a purple sphere, and choline as sticks and colored by heteroatom. The rectangle in dashed lines highlight the location in the structure of the enlarged view of $Na^+$ (c). **c)** Close-up view of the $Na^+$ cation, with interacting residues depicted in sticks and the water molecule as a sphere.

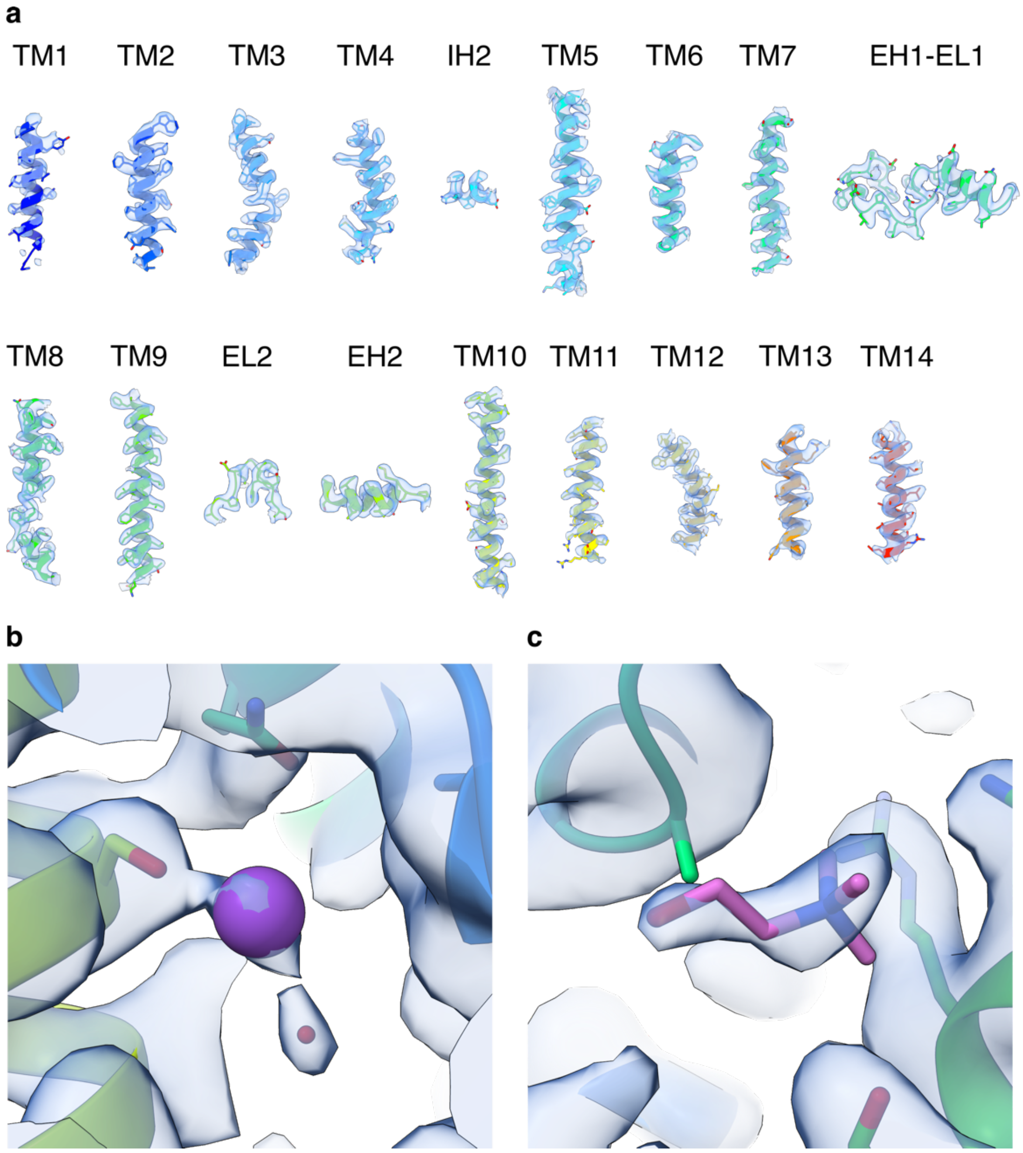


Supplementary Figure 11. **Cryo-EM map and model-to-map fit of choline-bound sfCHT.** The map threshold was set to an RMSD threshold of 6.30 Å in panel (a), and to 4.10 Å in (b) and (c). **a)** The different segments of sfCHT, as named as in Supplementary Figure 1, are depicted in cartoon format with the side chains shown in sticks. The cryo-EM density map is displayed around these segments. **b)** Close-up view of the $Na^+$ cation shown as spheres, interacting with residues depicted in sticks and the water molecule as a sphere. **c)** Close-up view of the ligand choline bound near the intracellular side of the protein and the interacting ligands.

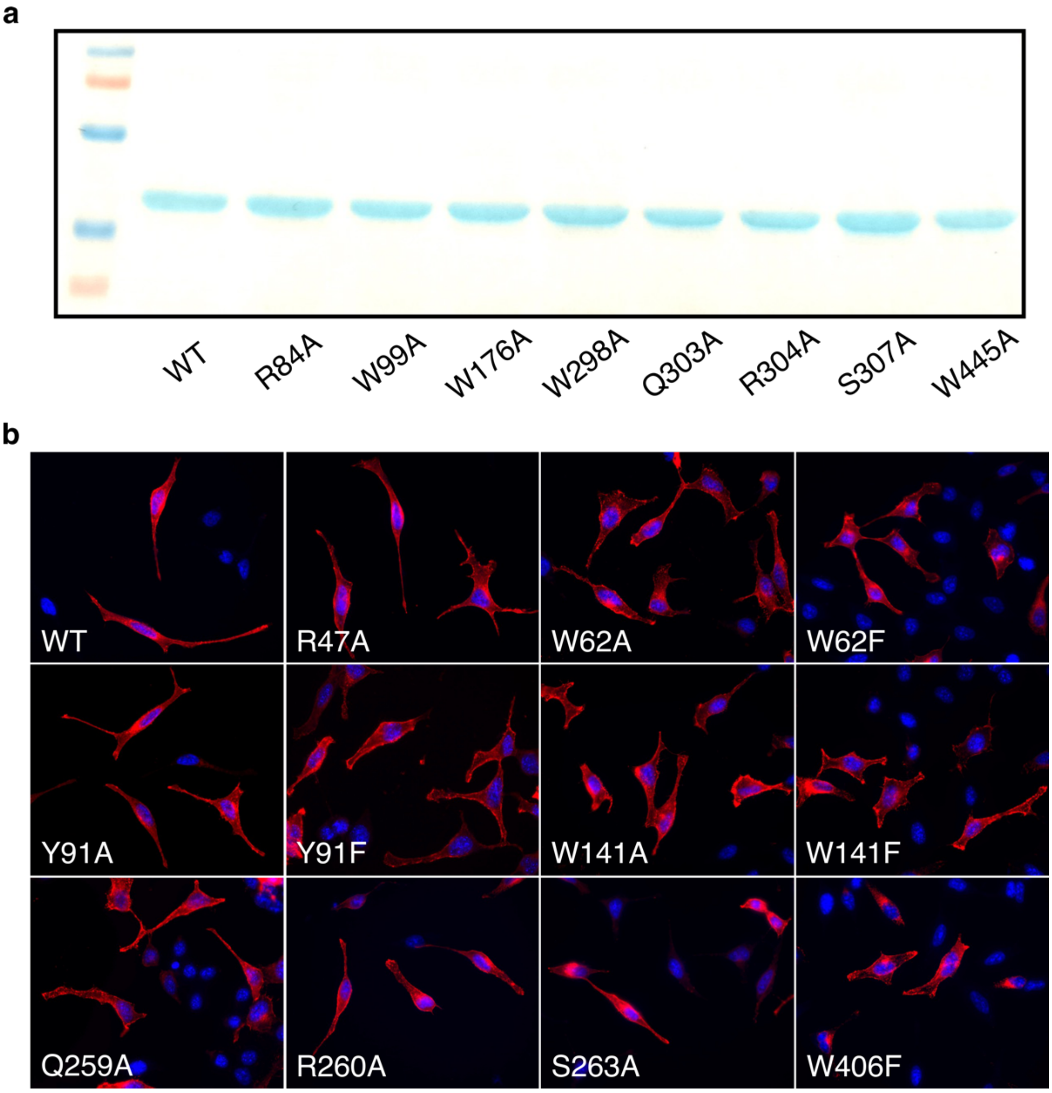


Supplementary Figure 12. **SDS-PAGE analysis of liposome reconstituted WT and variant sfCHT (a) and expression of WT and variant CHT1 expressed in HeLa cells (b).** Cells were seeded on 24-well plates, transfected and assayed 24 h post-transfection. Representative images of immunofluorescence of WT and variant CHT1. CHT1 (red) and Hoechst 33342 (blue) labelling is shown. All CHT1 variants were expressed at a similar level and reached the plasma membrane as WT CHT1.

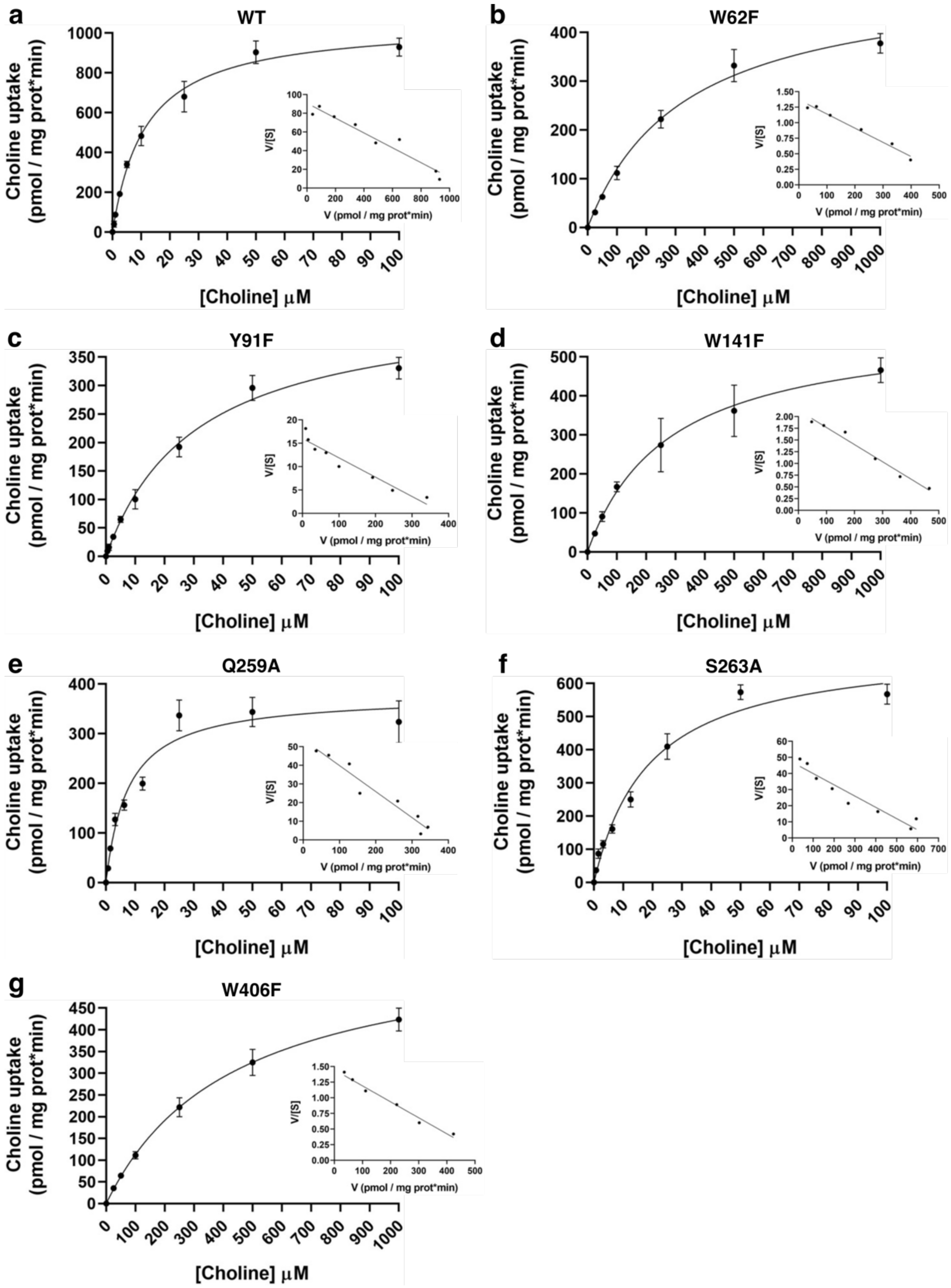


Supplementary Figure 13. **WT and variant CHT1-induced uptake of [$^3$H]-choline kinetic analysis.** Data (mean±SD) from representative experiments run in triplicates are shown. Inset: Eadie-Hofstee transformation.

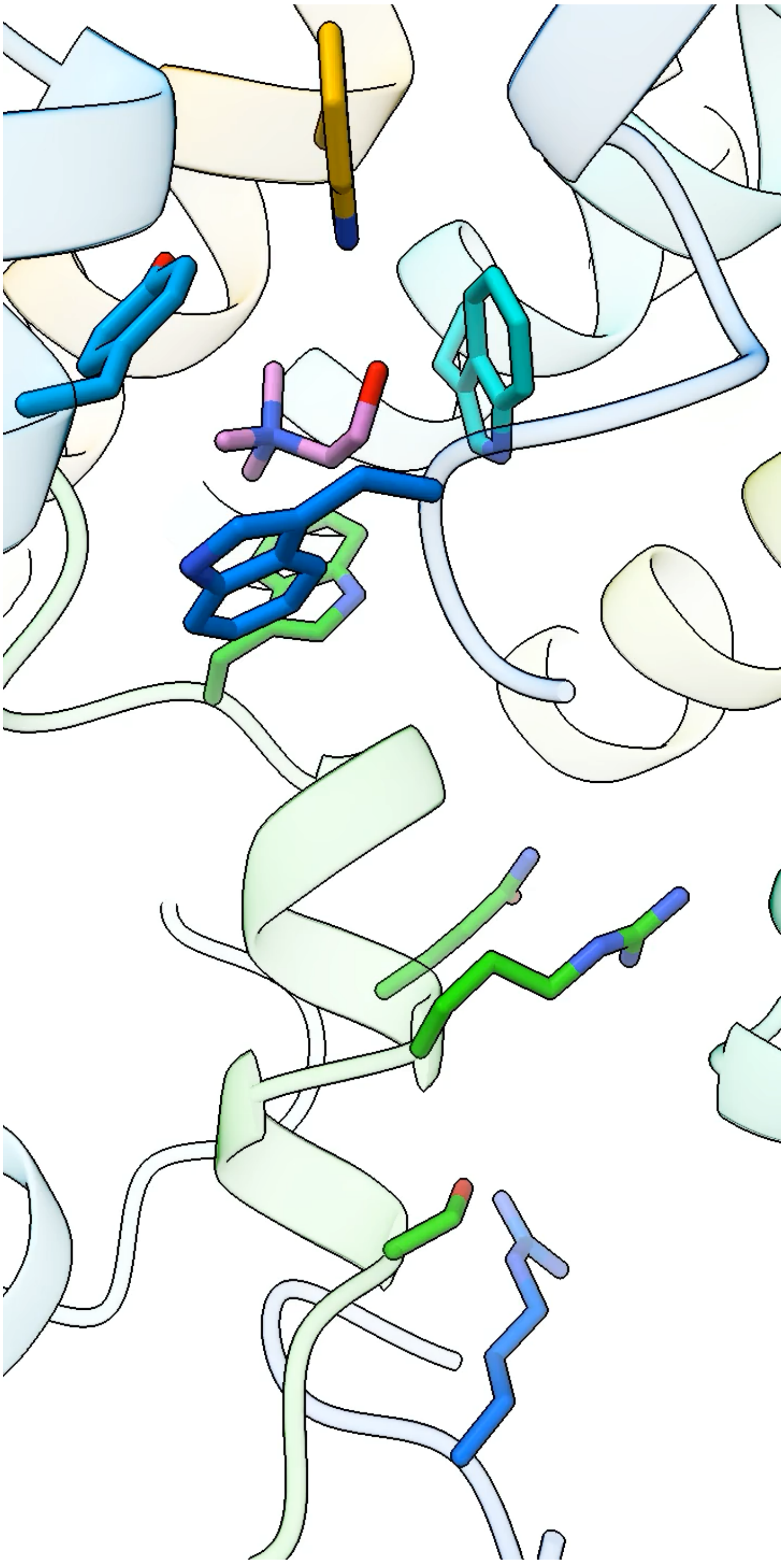

Supplementary Movie 1. **Choline translocation from the substrate binding site to the cytoplasmic vestibule.** Morph with PELE predicted poses and the cryo-EM structure in the presence of choline.
